# Aberrant N-glycosylation is a therapeutic target in carriers of a common and highly pleiotropic mutation in the manganese transporter ZIP8

**DOI:** 10.1101/2024.06.28.601207

**Authors:** Vartika Tomar, John Kang, Ruxian Lin, Steven R. Brant, Mark Lazarev, Caitlin Tressler, Kristine Glunde, Natasha Zachara, Joanna Melia

## Abstract

The treatment of defective glycosylation in clinical practice has been limited to patients with rare and severe phenotypes associated with congenital disorders of glycosylation (CDG). Carried by approximately 5% of the human population, the discovery of the highly pleiotropic, missense mutation in a manganese transporter ZIP8 has exposed under-appreciated roles for Mn homeostasis and aberrant Mn-dependent glycosyltransferases activity leading to defective N-glycosylation in complex human diseases. Here, we test the hypothesis that aberrant N-glycosylation contributes to disease pathogenesis of ZIP8 A391T-associated Crohn’s disease. Analysis of N-glycan branching in intestinal biopsies demonstrates perturbation in active Crohn’s disease and a genotype-dependent effect characterized by increased truncated N-glycans. A mouse model of ZIP8 391-Thr recapitulates the intestinal glycophenotype of patients carrying mutations in ZIP8. Borrowing from therapeutic strategies employed in the treatment of patients with CDGs, oral monosaccharide therapy with N-acetylglucosamine ameliorates the epithelial N-glycan defect, bile acid dyshomeostasis, intestinal permeability, and susceptibility to chemical-induced colitis in a mouse model of ZIP8 391-Thr. Together, these data support ZIP8 391-Thr alters N-glycosylation to contribute to disease pathogenesis, challenging the clinical paradigm that CDGs are limited to patients with rare diseases. Critically, the defect in glycosylation can be targeted with monosaccharide supplementation, providing an opportunity for genotype-driven, personalized medicine.

## Main

Approximately 5% of the human population carries a missense mutation in the gene *SLC39A8* encoding the metal transporter protein ZIP8 (rs13107325; ZIP8 A391T).^1^ ZIP8 regulates systemic manganese homeostasis.^2, 3^ ZIP8 A391T ranks in the top 10 most pleiotropic variants in the human genome across 42 traits^4^ and is associated with many complex diseases, including an increased risk of Crohn’s disease, schizophrenia, obesity, scoliosis, dyslipidemia, pain, and susceptibility to Covid-19 infection, as well as a decreased risk of hypertension, alcohol misuse disorders, and Parkinson’s disease.^4–9^ These fundamental genetic observations highlight the role of Mn homeostasis in human health and disease.

Individuals with rare, biallelic loss-of-function mutations in *SLC39A8* present with undetectable Mn levels and defective glycosylation marked by impaired N-glycan branching and are classified as a type 2 congenital disorder of glycosylation (SLC39A8-CDG).^10^ ^11, 12^ As Mn is used by many glycosyltransferases to coordinate nucleotide sugars substrates, decreases in Mn reduce the efficiency of Mn-dependent glycosyltransferases, favoring the formation of less branched N-glycan structures.^10^ In contrast to the loss-of-function mutations, ZIP8 A391T is a hypomorphic mutation.^7, 13^ ZIP8 391-Thr results in a relative Mn insufficiency likely through impaired resorption of Mn from bile as well as impaired intestinal absorption.^2, 3^ Blood Mn levels are decreased by approximately 15% in heterozygous carriers and 30% in homozygous carriers.^7, 14, 15^ Despite the more modest reduction in Mn levels compared to those found in patients with SLC39A8-CDG, heterozygous and homozygous carriers of ZIP8 391-Thr also exhibit defective glycosylation with decreased levels of complex N-glycans (tri- and tetra-antennary species) and increased biantennary N-glycans in plasma.^14, 16^ Although CDGs have historically been considered rare in clinical practice, we hypothesize that ZIP8 A391T-related pathobiology represents an attenuated form of CDG with the potential to impact disease risk in a high number of individuals worldwide given the allele frequency and impact even with heterozygosity.^17–19^

To test this hypothesis, it is necessary to study how aberrant N-glycosylation may contribute to a ZIP8 391-Thr-associated disease, and if treatment targeting glycosylation can ameliorate disease. Previous studies have focused on the features of the plasma N-glycome in ZIP8 391-Thr carriers.^14, 16^ There are no data on the effect of ZIP8 391-Thr on N-glycosylation in tissues in humans, although animal studies have shown an effect on the tissue N-glycome in the brain.^20^ Importantly, ZIP8 391-Thr is also highly associated with variation in 22 brain structures measured through MRI-based studies performed through the UK Biobank.^21, 22^ These data implicate alterations to the N-glycome as a possible driver of structural – and by extension, physiologic activities of a disease-associated organ system. Membrane-bound and secreted proteins are heavily N-glycosylated with complex N-glycans, making study at mucosal interfaces particularly important.^23, 24^ Given the association between ZIP8 391-Thr and risk of Crohn’s disease, here we examine accessible human tissues from intestinal biopsies relevant to ZIP8 391-Thr-associated Crohn’s disease and also examine an animal model of inflammatory bowel disease to test the hypothesis that ZIP8 391-Thr-associated disease is at least in part a form of CDG that is amenable to treatment as a disease of disordered glycosylation.

## Results

### The ileal epithelium is dominated by L-PHA-positive complex N-glycans in health and truncated sWGA-positive N-glycans in active Crohn’s disease

We began by asking if all patients with Crohn’s disease, regardless of genotype, exhibit evidence of aberrant N-glycosylation in diseased tissue.^25^ We focused on the ileum as the genetic association between ZIP8 391-Thr and Crohn’s disease was observed to be significantly greater in patients with ileal Crohn’s disease vs. non-ileal Crohn’s disease location.^26^ We chose two lectins to delineate between complex N-glycans (L-PHA) and truncated N-glycans that terminate in N-acetylglucosamine (succinylated WGA [sWGA]) (**Fig. 1A**).^27^ In healthy individuals, the ileal epithelial compartment is dominated by L-PHA stained cells with limited sWGA staining. In contrast, in active Crohn’s disease, there is increased sWGA staining. We used a ratio of sWGA/L-PHA intensities normalized to DAPI to quantify the difference in staining pattern; sWGA/L-PHA ratio of intensities increased from mean 0.69 to 1.65 in active disease (p = <0.001) (**Fig. 1B**). The increased abundance of truncated N-glycans is in congruence with transcriptional down-regulation of branching glycosyltransferases with active Crohn’s disease: We studied differential expression of glycosyltransferases of the N-glycosylation cascade in patients with active Crohn’s disease using a publicly-available bulk RNAseq (**Fig. 1C; Supplementary Table 1).**^28^ This dataset is derived from ileal biopsies from pediatric patients with newly diagnosed Crohn’s disease prior to initiation of medications as part of an inception cohort study.^28, 29^ The most abundant glycosyltransferase in the ileum by transcript is Mn-dependent *MGAT4B*, and *MGAT4B* is highly differentially expressed with down-regulation in active disease compared to healthy individuals (q = <0.000001). The Mn-utilizing *MGAT3* (q = <0.000001), *MGAT4A* (q = 0.000011), and *MGAT5* (q = <0.000001) are also down-regulated, while there is up-regulation of most capping and extension enzymes (e.g. *B3GNT9* (q = <0.000001), *GCNT2* (q = <0.000001), consistent with compensatory or ‘self-correction’ mechanisms that are activated when branching is impaired.^30, 31^ Notably, many of the capping and extension enzymes also require Mn.^11^

**Fig. 1.**
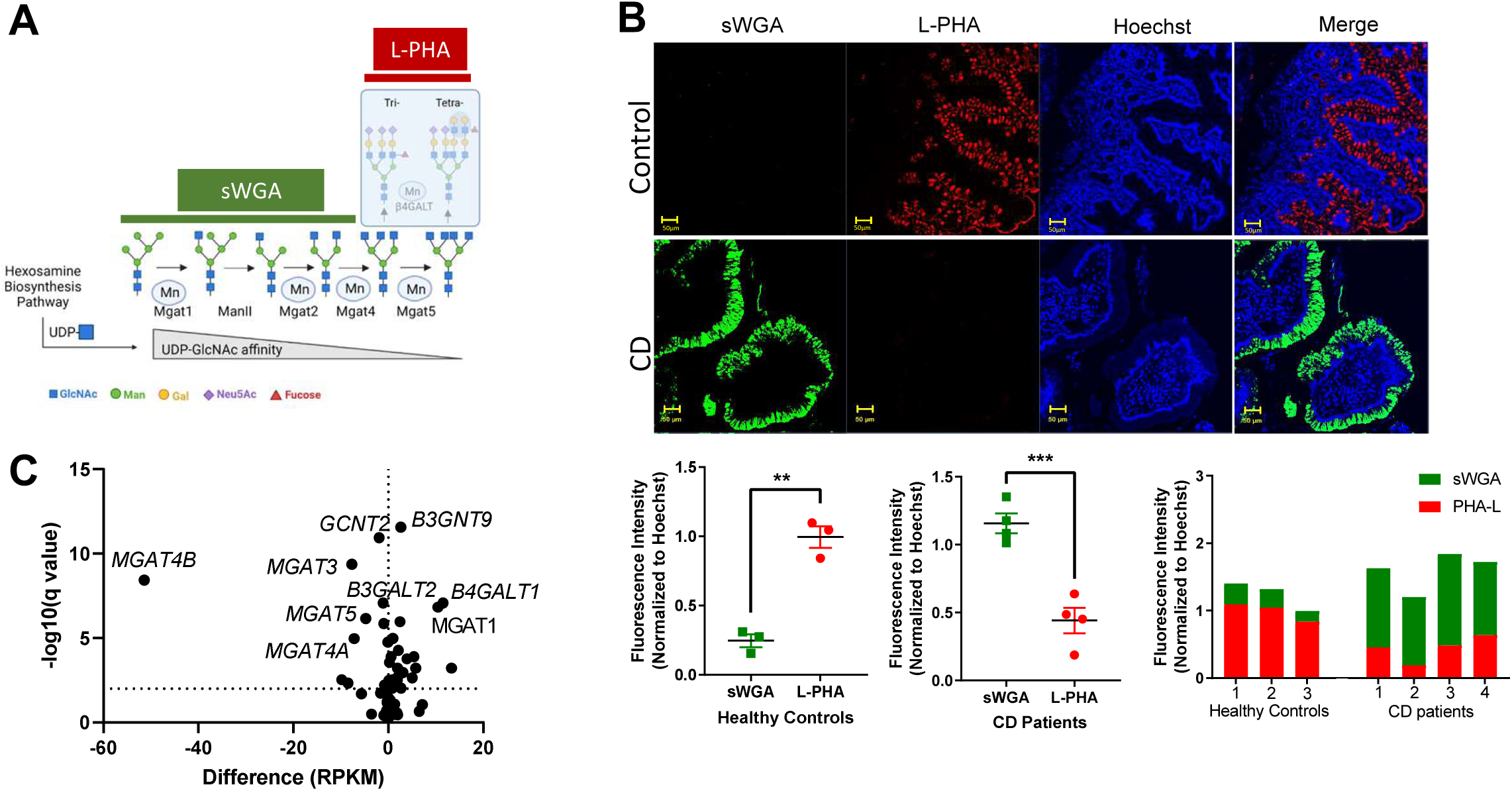
The ileal epithelium is dominated by L-PHA-positive, complex N-glycans in health and truncated, sWGA-positive, N-glycans in active Crohn’s disease. (**A**) Schematic of N-glycan branching cascade demonstrating the glycan targets of lectins and manganese (Mn) utilization. sWGA (green) stains terminal GlcNAc residues. L-PHA (red) stains complex tri- and tetra-antennary N-glycans. Relative UDP-GlcNAc affinity of glycosyltransferases is also represented (300-fold change from Mgat1 to Mgat5). Figure generated in Biorender, informed by Cummings, et al (Ref. 26). (**B**) Confocal laser-scanning triple-label immunofluorescence microscopy images of human ileal tissues paraffin sections, stained for PHA-L (red), sWGA (green), Hoechst (blue) and merged image. Samples were incubated with Fluorescein dyes (PHA-L 639, sWGA 488 and Hoechst 405) 10 µg/ml in blocking buffer for 1 hour at room temperature. Scale bar is 50µm. n = 3-4. Fluorescence intensity of sWGA and L-PHA normalized to Hoechst was measured using Metamorph to compare healthy individuals and those with Crohn’s disease (CD). Ratio of sWGA/L-PHA also depicted. Individual data points, mean, and SEM graphed with statistical significance are determined by t-test with p-value indicated by two asterisks (<.01) or three asterisks (<.001). (**C**) Volcano plot of glycosyltransferase genes differentially expressed in ileal mucosal biopsies from male and female pediatric patients with severe ileal Crohn’s disease compared to non-IBD controls. This is a secondary analysis of RNAseq data GSE57945. n=42 controls, n=63 CD patients. There is an enrichment for MGAT-related pathway members with increased expression of *MGAT1* and decreased expression of *MGAT3*, *MGAT4A/4B*, and *MGAT5*. *MGAT4B* is the most highly expressed gene (Supplemental Table 1).

### Truncated sWGA-positive N-glycans are increased at the apical/brush border of ileal epithelial cells in ZIP8 391-Thr carriers with Crohn’s disease

If active Crohn’s disease is characterized by a loss of complex N-glycans in the ileal epithelial compartment, we hypothesized that ZIP8 391-Thr carriers would be even more susceptible to impaired N-glycan branching if there is transcriptional down-regulation of glycosyltransferases and relative Mn insufficiency limiting glycosyltransferase activity. When we stratify by ZIP8 genotype in a cohort of patients with ileal Crohn’s disease, measured intensities of sWGA or L-PHA or sWGA/L-PHA ratio in ileal biopsies with or without active inflammation did not differ by genotype, however, localization was significantly different. Carriers of ZIP8 391-Thr showed enhanced apical/brush border staining by sWGA (p=0.01) and increased apical overlap of sWGA and L-PHA (p=0.028) (**Fig. 2A,B**). Together, these data support that ZIP8 391-Thr is associated with aberrant N-glycosylation with an increase in truncated N-glycans (stained by sWGA) at the apical/brush border of enterocytes in the ileum in patients with active Crohn’s disease. These data align with the N-glycome signature in plasma where tri- and tetra-antennary N-glycans are decreased in ZIP8 391-Thr carriers with Crohn’s disease.^16^

**Fig. 2.**
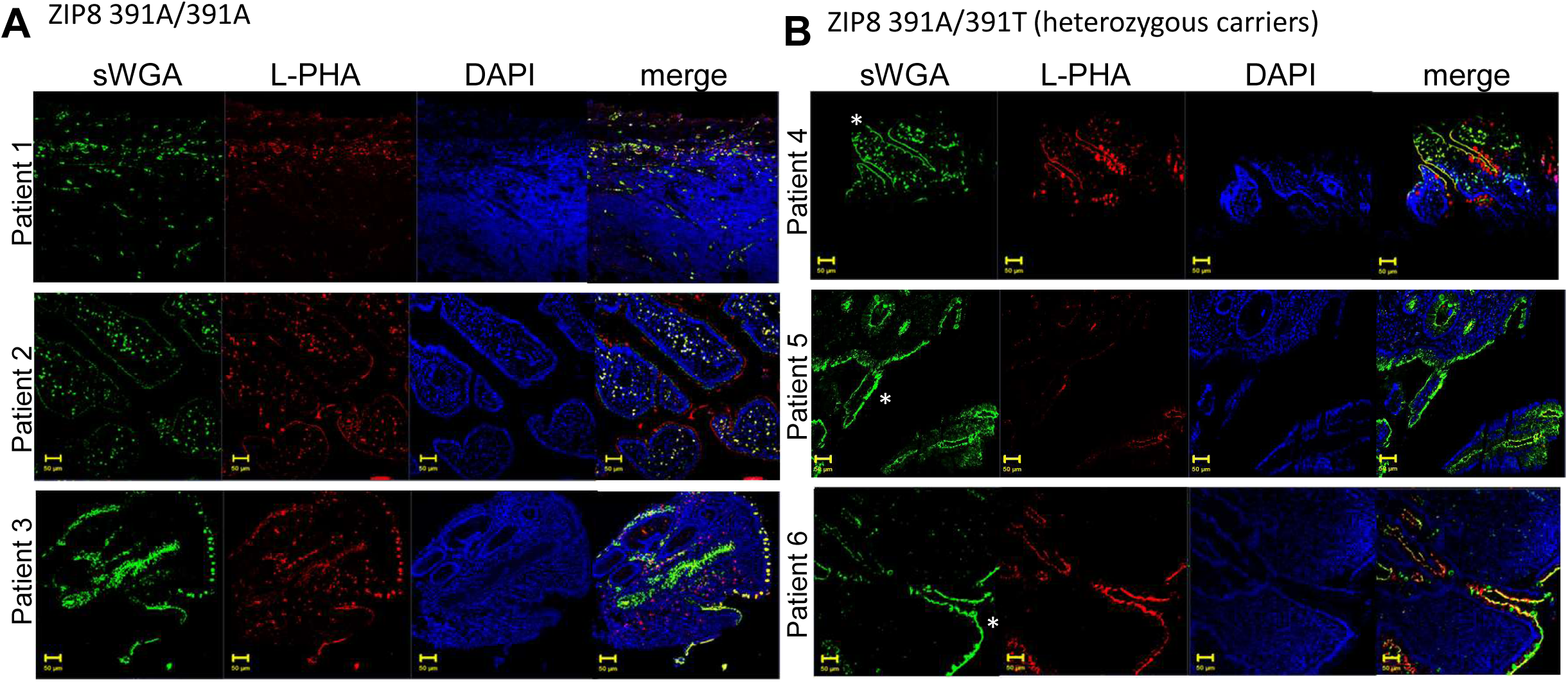
Truncated sWGA-positive N-glycans are increased at the apical/brush border of ileal epithelial cells in ZIP8 391-Thr carriers with Crohn’s disease. Representative lectin immunofluorescence images of ileal biopsies of genotyped patients with active Crohn’s ileitis in (**A**) ZIP8 391A/391A (non-carriers) and (**B**) ZIP8 391A/391T (ZIP8 391-Thr heterozygous carriers). Confocal laser-scanning triple-label immunofluorescence microscopy images of human ileal tissues paraffin sections, stained for L-PHA (red), sWGA (green), Hoechst (blue) and merged image. Samples were incubated with Fluorescein dyes (L-PHA 639, sWGA 488 and Hoechst 405) 10 µg/ml in blocking buffer for 1 hour at room temperature. Scale bar: 50µm. Asterisk highlights enhanced localization of sWGA-staining to the apical membrane/glycocalyx of epithelial cells. To focus on the epithelial compartment, sWGA and L-PHA distribution and overlap were blindly scored by two investigators. Representative images from n=3 patients/genotype included in figure; histology reviewed and scored for n= 9 ZIP8 391A/391A; n= 6 ZIP8 391A/391T individuals.

### ZIP8 391-Thr-associated defect in N-glycosylation in the ileal epithelial compartment is recapitulated in Zip8 393T-knock-in mice

Our next question was if a Zip8 genotype-driven effect on N-glycosylation in the ileum could be recapitulated in a mouse model of ZIP8 391-Thr. We have previously generated a Zip8 393T-knock-in mice (murine Zip8 393T-KI is equivalent to human ZIP8 391T) using CRISPR-Cas9.^16^ Ileal sections were taken from Zip8^+/+^, Zip8^+/393T^ and Zip8^393T/393T^ mice and lectin staining was performed for sWGA and L-PHA as we performed for the human tissues (**Fig. 3A,B**). Similar to the human studies, we observed a significant effect of the Zip8 genotype on N-glycan branching. L-PHA was the dominant lectin in the Zip8^+/+^ mice with a near absence of sWGA staining, much like the dominant pattern observed in ileal sections from healthy individuals (**Fig. 1B**). In contrast, there was enhanced sWGA staining in the crypts and along the brush border in Zip8^+/393T^ and Zip8^393T/393T^ animals. There is L-PHA staining in the crypts and decreased abundance along the brush border in Zip8^+/393T^ and Zip8^393T/393T^ animals compared to Zip8^+/+^. As a complementary method, we used matrix-associated laser desorption/ionization (MALDI) mass spectrometry imaging (MSI) following on-tissue PNGase digest to measure the differential abundance of N-glycan species in Zip8^+/+^ and Zip8^393T/393T^ mice (**Fig. 3C; Supplementary Fig. 1**). This analysis demonstrated decreased abundance of complex, branched species in Zip8^393T/393T^ mice.

**Fig. 3.**
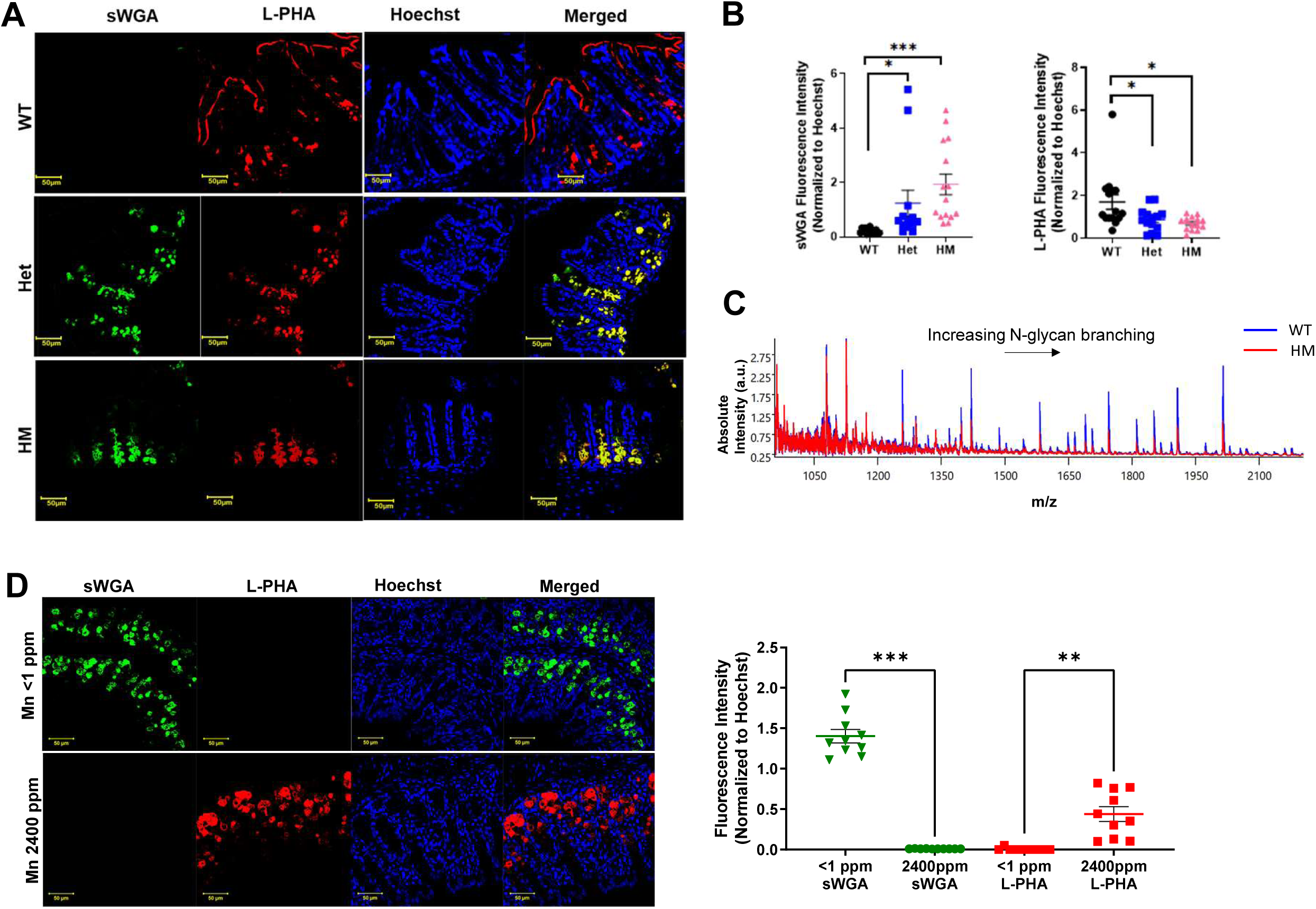
ZIP8 391-Thr-associated defect in N-glycosylation in the ileal epithelial compartment is recapitulated in Zip8 393T-knock-in mice. (**A**) Confocal laser-scanning triple-label immunofluorescence microscopy images of Zip8^+/+^ (WT), Zip8^+/393T^ (Het) and Zip8^393T/393T^ (HM) ileal tissues paraffin sections, stained for L-PHA (red), sWGA (green), Hoechst (blue) and merged image. Samples were incubated with Fluorescein dyes (L-PHA 639, sWGA 488 and Hoechst 405) 10 µg/ml in blocking buffer for 1 hour at room temperature. Scale bar: 50µm. N=5 male and female mice/genotype with n=3-5 fields of view/mice imaged. (**B**) Quantification of sWGA and L-PHA fluorescence intensity normalized to Hoechst measured using Metamorph. Statistical significance determined by one-way ANOVA, p-value indicated by one asterisk (<.05) or three asterisks (<.001). (**C**) Averaged spectra of matrix-associated laser desorption/ionization (MALDI) mass spectrometry imaging (MSI) following on-tissue PNGase F digest to measure the differential abundance of N-glycan species in transverse section of distal ileal tissue of Zip8^+/+^ and Zip8^393T/393T^ mice. Higher m/z species represent tri- and tetra-antennary N-glycan branching. N=4 male mice/genotype. (**D**) Confocal laser-scanning triple-label immunofluorescence microscopy images of Jackson C57BL/6 male mice fed purified diet containing variable Mn (<1ppm = Mn deficient and 2400ppm = Mn excess) for 4 weeks. Ileal tissues paraffin sections, stained for L-PHA (red), sWGA (green), Hoechst (blue) and merged image. N = 3 male mice in each group, 2-3 fields of view imaged and quantified per mouse. Scale bar: 50µm. Individual data points, mean, and SEM graphed with statistical significance determined by one-way ANOVA, p-value indicated by two asterisk (<.01) or three asterisks (<.001).

To determine the dependence of ileal epithelial N-glycosylation glycophenotype on dietary Mn and host Mn status, we fed C57BL/6 WT mice purified diets with variable Mn content: Mn <1 ppm (Mn deficient) or 2400 ppm (Mn excess) (**Fig. 3D; Supplementary Fig. 2**). The glycophenotype of the mice fed the Mn deficient diet phenocopy Zip8^393T/393T^ animals with enhanced abundance of sWGA staining (p = 0.0002). In contrast, L-PHA was the dominant lectin in the animals fed Mn in excess (p < 0.0011). Together, these data are consistent with a genotype effect of Zip8 and an effect of host Mn status (here determined by dietary Mn intake) on the N-glycan branching of the ileal epithelial compartment and position the animal model to test therapeutic strategies.

### Oral GlcNAc supplementation restores complex N-glycan branching in intestinal epithelial cells in Zip8^+/393T^ and Zip8^393T/393T^ mice

The rate-limiting substrate for N-glycan branching is N-acetylglucosamine (GlcNAc) complexed to uridine diphosphate (UDP). There is a 300-fold decrease in affinity for UDP-GlcNAc moving from MGAT1 to MGAT5 (**Fig. 1**).^32, 33^ Building on studies demonstrating that N-glycan branching can be rescued by GlcNAc supplementation, we hypothesized that GlcNAc supplementation would improve the efficiency of the N-glycan cascade despite Mn insufficiency.^1, 34^ This approach is adapted from the therapeutic strategy employed to treat patients with CDGs with loss-of-function mutations in glycosyltransferases where excess monosaccharide supplementation can bypass enzyme defects.^35^ Glycan-targeting therapy spares the risks of a narrow therapeutic window of excess Mn supplementation given the association of excess Mn accumulation in the brain and Parkinsonian-like disease; further, impaired function of ZIP8 disrupts intestinal Mn absorption.^3, 36^ Glycan-targeting therapy also capitalizes on favorable safety and efficacy data of GlcNAc supplementation in patients with inflammatory bowel disease and other immune-mediated disorders, like multiple sclerosis.^37–40^

To test in vivo, we supplemented Zip8^+/+^, Zip8^+/393T^ and Zip8^393T/393T^ with GlcNAc 0.25 mg/ml in drinking water for 7 days and studied the change in N-glycosylation of the ileal epithelial compartment by lectin immunofluorescence (**Fig. 4A,B,C**). The GlcNAc-treated Zip8^+/393T^ and Zip8^393T/393T^ animals show enhanced L-PHA staining, consistent with increased tri- and tetra-antennary N-glycan branching of the ileal epithelial cells and along the epithelial brush border with a reciprocal decrease in sWGA staining compared to sham. GlcNAc treatment reversed the sWGA/L-PHA ratio in the Zip8^+/393T^ mice (**Fig. 4B**), making L-PHA the dominant staining lectin and recapitulating the Zip8^+/+^ glycophenotype. The sWGA/L-PHA ratio does not fully reverse in the Zip8^393T/393T^ mice, but it is shifted towards the Zip8^+/+^ glycophenotype (**Fig. 4C**).

**Fig. 4.**
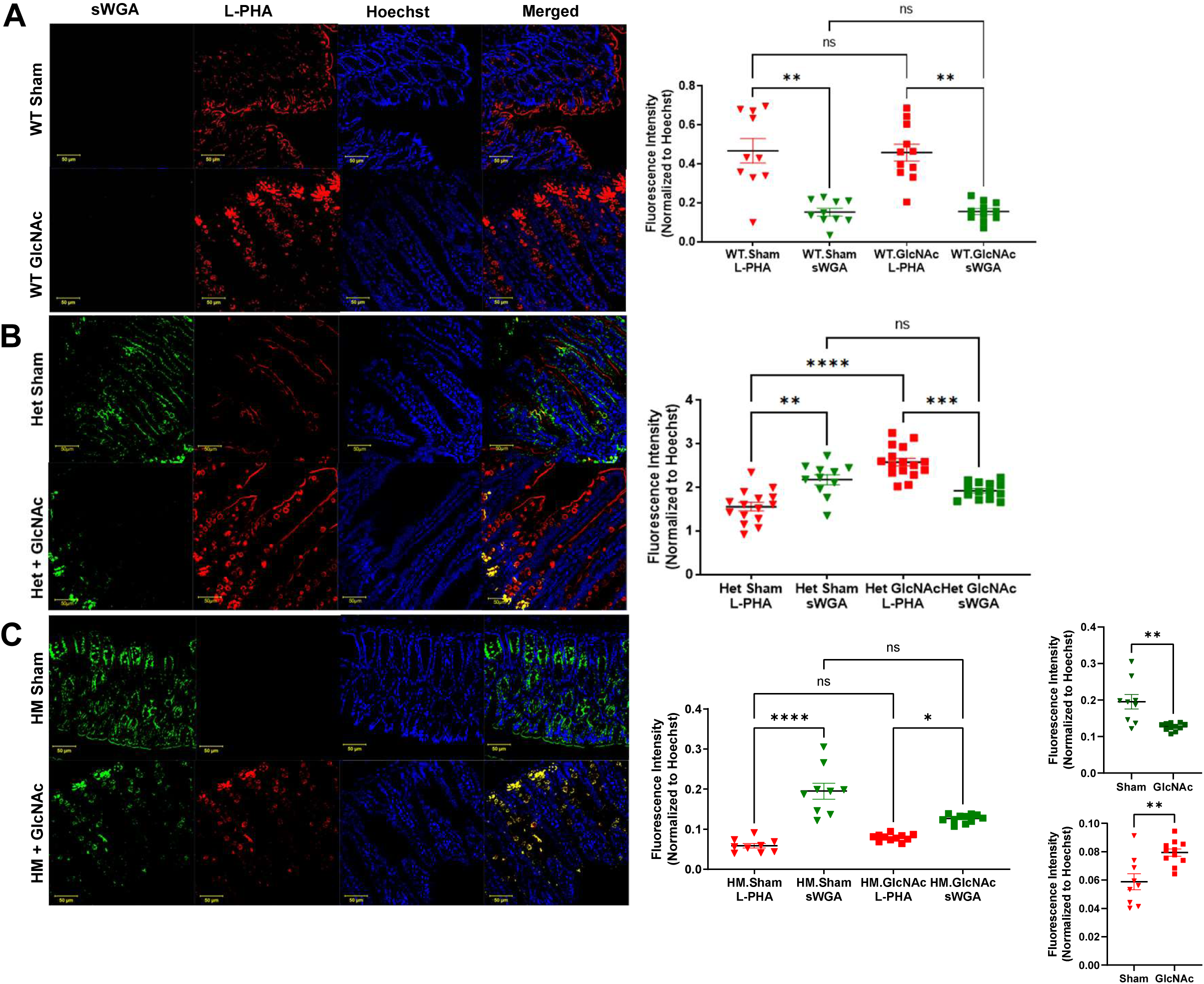
Oral GlcNAc supplementation restores complex N-glycan branching in intestinal epithelial cells in Zip8^+/393T^ and Zip8^393T/393T^ mice. Confocal laser-scanning triple-label immunofluorescence microscopy images of Zip8^+/+^ (WT) (**A**), Zip8^+/393T^ (Het) (**B**) and Zip8^393T/393T^ (HM) (**C**) mice ileal tissue paraffin sections, stained for sWGA (green), L-PHA (red), Hoechst (blue) and merged. Samples were incubated with Fluorescein dyes (L-PHA 639, sWGA 488, and Hoechst 405) 10 µg/ml in blocking buffer for 1 hour at room temperature. L-PHA and sWGA fluorescence intensity measured using Metamorph. Scale bar: 50µm. N=4-5 male and female mice/genotype with n=3-7 fields of view/mice imaged. Individual data points, mean, and SEM are graphed. Statistical significance was determined by Kruskal-Wallis one-way ANOVA with Dunn’s multiple comparisons testing; inset graphs in (C) show statistics from Mann-Whitney t-test that show significant difference despite non-significance by ANOVA; p-value indicated by one asterisk (<.05), two asterisk (<.01), three asterisks (<.001), or four asterisks (<.0001).

### Defects in bile acid homeostasis and intestinal permeability are ameliorated with oral GlcNAc supplementation in Zip8^+/393T^ and Zip8^393T/393T^ mice

Regulation of bile acid homeostasis through the FXR-Fgf15 pathway is a primary role of the ileal epithelium, and uptake of bile acids has been shown to be dependent on N-glycosylation.^41^ We have previously shown that carriers of ZIP8 391-Thr with Crohn’s disease have a distinct ileal and rectal mucosal microbiota profile associated with aberrant bile acid homeostasis.^42^ Further, in a subset of patients with ileocolonic Crohn’s disease, ZIP8 391-Thr carriers had decreased circulating FGF19 levels.^42^ These associations in patient cohorts between ZIP8 genotype and bile acid homeostasis were recapitulated in the Zip8 393T-KI mouse model where total bile acids are increased in the liver and stool, consistent with increased production of bile secondary to decreased re-uptake of bile acids in the ileum and loss of FXR-Fgf15 signaling. We therefore tested if GlcNAc supplementation would ameliorate defects in bile acid homeostasis in Zip8^+/393T^ and Zip8^393T/393T^ mice as functional evidence of amelioration epithelial physiology. Consistent with this hypothesis, GlcNAc supplementation improved the defect in bile acid homeostasis, decreasing total bile acids in stool (**Fig. 5A**) with increased FXR-Fgf15 signaling as measured by *Fgf15* mRNA expression in ileum (**Fig. 5B**) and *Cyp7a1* mRNA expression in liver (**Fig. 5C**). As a second measure of intestinal physiology, we studied soluble CD14 (sCD14), a blood-based marker of gut barrier function.^43^ At baseline, sCD14 is increased in Zip8^+/393T^ and Zip8^393T/393T^ compared to Zip8^+/+^ mice. These data are consistent with a prior report in another strain of Zip8 393T-KI mice.^44^ Oral GlcNAc supplementation improved sCD14 towards Zip8^+/+^ levels, although sCD14 remained elevated in Zip8^+/393T^ and Zip8^393T/393T^ (**Fig. 5D**).^45^

**Fig. 5.**
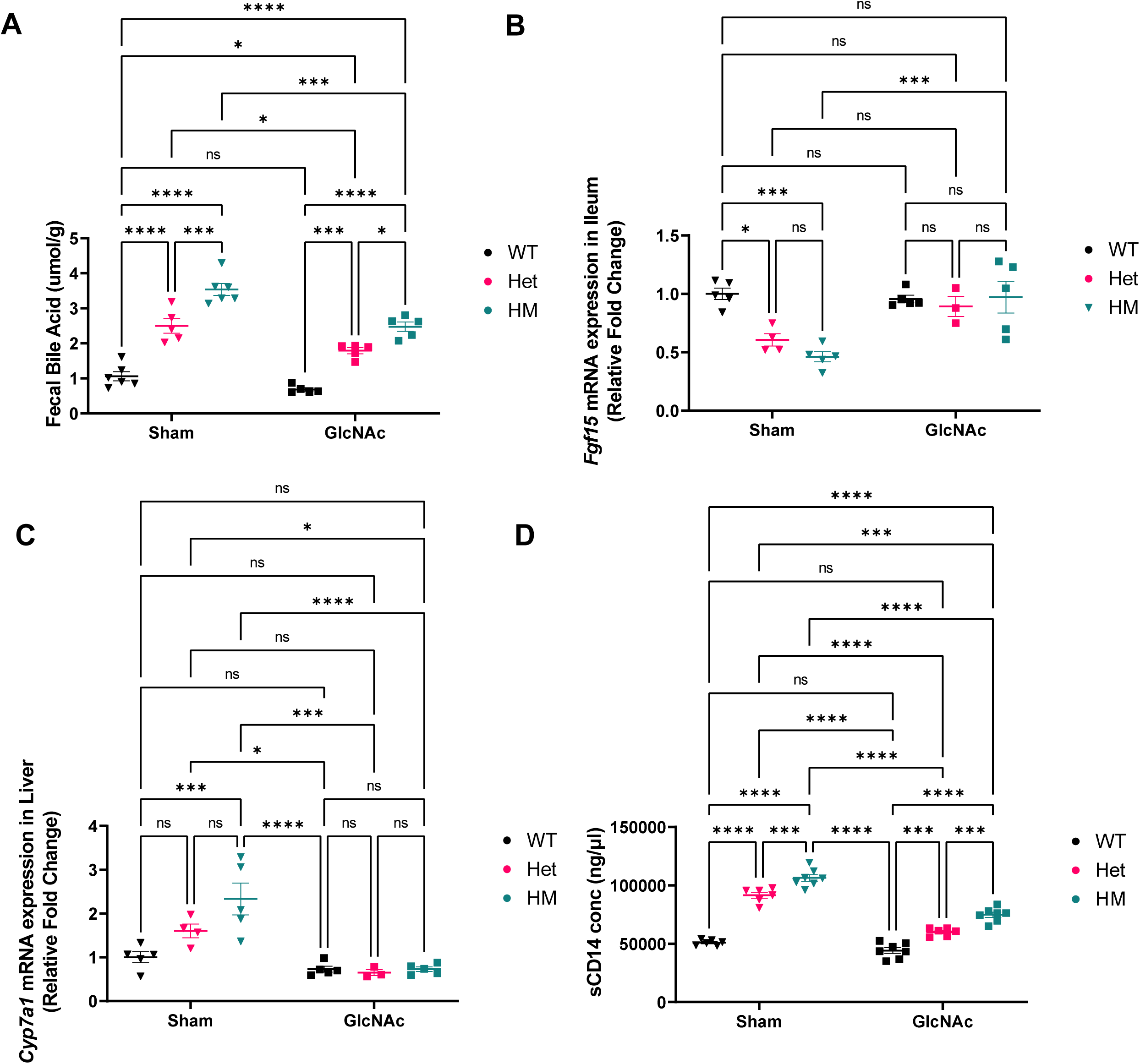
Defects in bile acid homeostasis and intestinal permeability are ameliorated with oral GlcNAc supplementation in Zip8^+/393T^ and Zip8^393T/393T^ mice. (**A**) Total fecal bile acid in Zip8^+/+^ (WT), Zip8^+/393T^ (Het), and Zip8^393T/393T^ (HM) mice with and without GlcNAc supplementation. (**B**) mRNA expression of *Fgf15* in ileum and (**C**) *Cyp7a1* in liver in Zip8^+/+^ (WT), Zip8^+/393T^ (Het), and Zip8^393T/393T^ (HM) mice with and without GlcNAc supplementation. mRNA data interpolated from standard curve using four parameter logistic (4-PL) curve (GraphPad Prism 9). (**D**) Serum sCD14 concentration using mouse CD14 Immunoassay Kit (#MC140). Data are represented as mean ± SEM. Statistical significance was determined by two-way ANOVA with multiple comparison testing. p-value indicated by one asterisk (<.05), two asterisk (<.01), three asterisks (<.001), or four asterisks (<.0001). n=5-6 male mice/genotype.

### Oral GlcNAc supplementation rescues colitis phenotype in Zip8^393T/393T^ mice

Finally, we^16^ and others^44^ have shown that Zip8^+/393T^ and Zip8^393T/393T^ mice have delayed recovery after chemical-induced colitis, a common model of intestinal injury and repair in studies of inflammatory bowel disease.^46^ To test if GlcNAc would ameliorate this phenotype, Zip8^+/+^ and Zip8^393T/393T^ mice were pre-treated with GlcNAc 0.25mg/ml in the drinking water for 7 days, exposed to 2% dextran sodium sulfate (DSS) for 5 days, and allowed to recover to 14 days with continued GlcNAc supplementation (**Fig. 6A**). GlcNAc rescued the Zip8^393T/393T^ mice with early recovery of body weight (**Fig. 6B**), decreased disease activity index (**Fig. 6C**), improvement in colon length (**Fig. 6D**), cecal weight (**Fig. 6E**), and spleen size (**Fig. 6F**), and improvement in colon histology (**Fig. 6G**). GlcNAc had no differential effect in Zip8^+/+^ mice, supporting a genotype-specific effect of GlcNAc supplementation in Zip8^393T/393T^ mice.

**Fig. 6.**
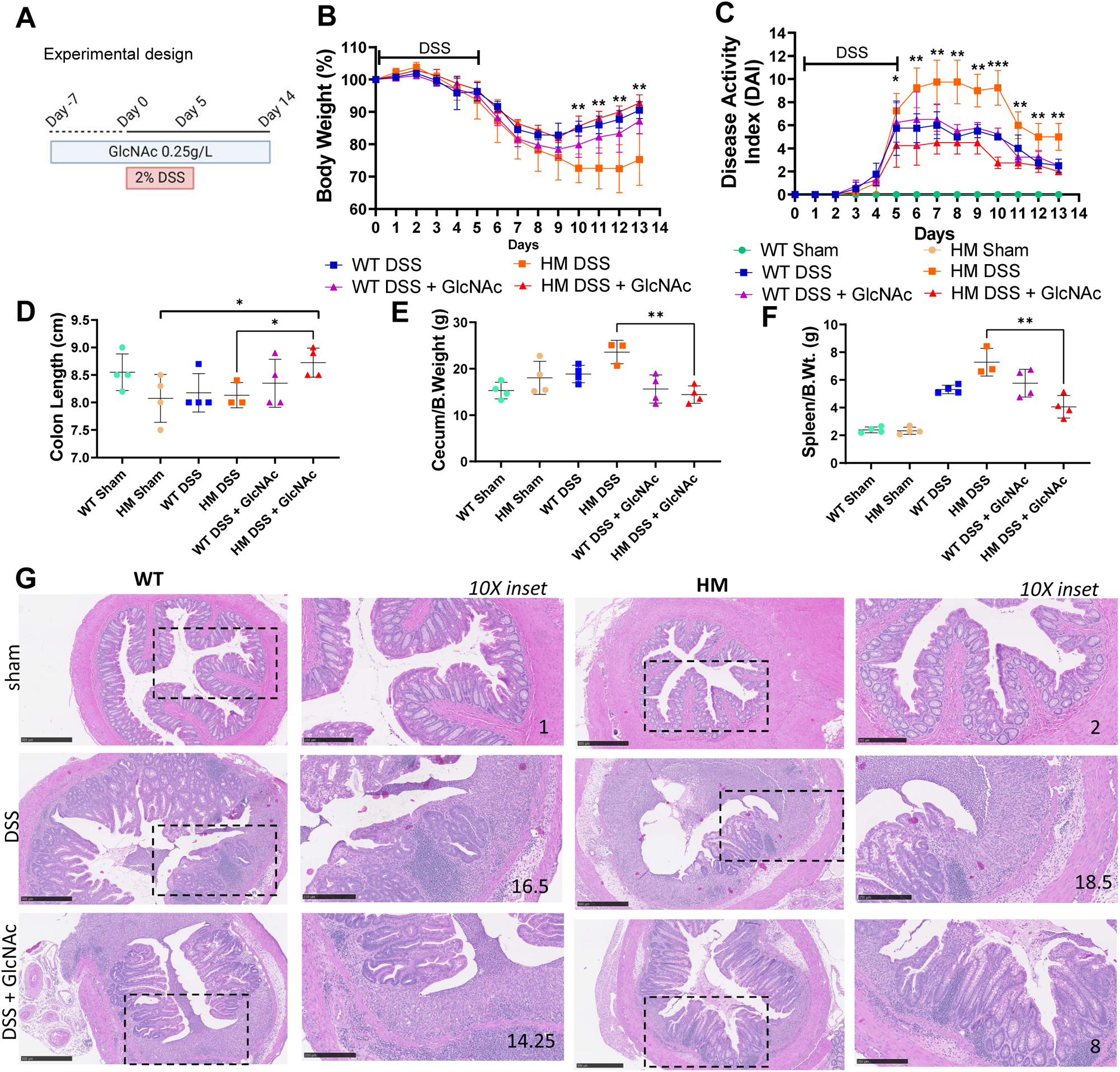
Oral GlcNAc supplementation rescues colitis phenotype in Zip8^393T/393T^ mice. Experimental schema (A). Effect of oral GlcNAc supplementation on body weight (B.Wt) loss (%) (**B**), disease activity index (DAI) (**C**), colon length (**D**), cecum/B.Wt ratios (**E**), spleen/B.Wt ratios (**F**), representative histology at sacrifice (Day 14) (**G**) in mice challenged in the DSS-induced colitis model. Colitis was induced by administering 2% DSS in distilled water for 5 days followed by distilled water for 9 days. In the GlcNAc-treated animals, GlcNAc (0.25g/L) dissolved in 2% DSS or distilled water was also administered for 7 days prior to DSS exposure and throughout the experimental period. Colitis was evaluated using DAI (body-weight loss, stool consistency and bleeding). Data represent the mean ± SEM of 3-4 animals in each control group (circle), DSS (square) and GlcNAc groups (triangle) for Zip8^+/+^ (WT) and Zip8^393T/393T^ (HM) male mice. Values are compared between HM DSS and HM GlcNAc groups using one-sided t-test with p-value indicated by one asterisk (<.05), two asterisk (<.01) or three asterisks (<.001). Day 14 data are not included for (A) and (B) as date of sacrifice with overnight fast prior to sacrifice. For the histologic images, H&E stained colonic specimens were imaged at 5X (scale bar: 500 µm) and 10X inset (scale bar represents 250 µm) with average histologic score included on inset for n=2-4 mice/genotype/experimental group (Koelink, et al.).

## Discussion

In this study, we demonstrate that the common and highly pleiotropic missense mutation, ZIP8 391-Thr, drives defective N-glycosylation that can contribute to the pathogenesis of Crohn’s disease as a model ZIP8 391-Thr-associated disease. These data challenge the clinical paradigm that congenital disorders of glycosylation (CDG) are limited to individuals with rare diseases with severe phenotypes, and suggest that glycan-targeting therapy could have important and broader applications.^35^

CDGs represent a clinically and genetically heterogeneous group of more than 130 diseases, most often inherited in an autosomal recessive pattern, with an estimated prevalence ranging from 1/10,000 to 1/22,000 individuals.^47, 48^ SLC39A8-CDG presents a unique opportunity to integrate the clinical observations made through the study of patients with rare, loss-of-function variants in ZIP8 and the highly pleiotropic and relatively common (in European ancestry and Latin American populations) mutation of ZIP8 391-Thr.^17–19^

Patients with SLC39A8-CDG have variable Mn blood levels, ranging from below the level of detection to low normal. ZIP8 has been reported to transport metals other than Mn to include Zn, Fe, cadmium, but in general, the trace metal deficiencies with SLC39A8-CDG have been limited to Mn with a single patient reported to have low normal blood Zn levels.^17–19^ A spectrum of glycosylation defects are characterized in patients with SLC39A8-CDG; the largest MALDI-TOF MS glycomics analyses showed an increase in hypogalactosylated precursor monosialo-mono-galacto-biantennary glycan (A2G1S1 m/z 2227) and a decrease in bisected glycans generated by MGAT3.^19^ Although there is clinical heterogeneity in patients with SLC39A8-CDG, there is consistent evidence supporting the role ZIP8 plays in Mn homeostasis and the interaction between Mn homeostasis and glycosylation. This is further supported in population-based studies demonstrating the association between ZIP8 391-Thr and lower blood Mn levels and features of the plasma N-glycome characterized by a relative decrease in trisialylated, trigalactosylated and triantennary N-glycan traits (*n* = 2763 individuals, ZIP8 391-Thr MAF 0.08).^16^ The global Zip8 knockout is embryonically lethal; however, inducible Zip8 knockout animals parallel these human data with decreased Mn levels in blood and tissues and decreased glycosyltransferase activity.^2^

The multi-systemic nature of diseases of disordered glycosylation may underscore the pleiotropic associations of ZIP8 391-Thr. Patients with SLC39A8-CDG present with multi-systemic disease with skeletal abnormalities, seizure disorders, developmental delay, hearing loss, and structural brain abnormalities by MRI.^17, 18^ In some patients, mitochondrial dysfunction (under the diagnosis of Leigh Syndrome) has also been attributed to Mn deficiency and dysfunction of mitochondrial Mn-dependent enzymes, including Mn superoxide dismutase. Notably, MnSOD enzyme activity has been reported to maintain redox homeostasis at <1% activity in prokaryotes, suggesting that the relative Mn insufficiency, but not deficiency in the context of ZIP8 391-Thr may spare mitochondrial dysfunction.^49, 50^

Treatment of patients with SLC39A8-CDG has utilized monosaccharide supplementation and Mn supplementation with close serial MRI imaging to monitor for Mn accumulation in the brain. ^17–19^ In ZIP8 391-Thr carriers, given the attenuated deficit in Mn and the large number of individuals potentially affected, treatment of the glycosylation defect mitigates the risk of Mn accumulation in the brain. The approach of monosaccharide supplementation tested here targets the 300-fold decrease in the affinity for UDP-GlcNAc moving from MGAT1 to MGAT5 to increase enzyme efficiency despite Mn insufficiency.^32, 33^ This approach is informed by monosaccharide supplementation to bypass primary glycosyltransferase defects to treat patients with multiple forms of CDG.^35^

Through study of patients with Crohn’s disease who carry ZIP8 391-Thr and our animal model of Zip8 393T-KI, our data show how the N-glycosylation defect associated with ZIP8 391-Thr may be particularly relevant when N-glycosylation is perturbed by the disease state, like Crohn’s disease.^25^ Approximately 1 in 10 patients with Crohn’s disease carries ZIP8 391-Thr, rising to 1 in 4 patients with Crohn’s disease of Ashkenazi Jewish descent.^16, 51, 52^ ZIP8 A391T most strongly associates with stricturing and penetrating disease of the ileum, the subtype of Crohn’s disease that is the most resistant to our current approved therapies^53^, and the most likely reason for patients to require surgery for their disease.^54–56^ The severity of disease observed in patients with ZIP8 391-Thr may reflect a “two-hit” hypothesis where the disease process itself impairs N-glycosylation on top of the baseline impairment in N-glycosylation that cannot be adequately reversed. These data lead to our hypothesis that glycan-directed therapy is a critical adjunctive therapy for the treatment of patients who carry ZIP8 391-Thr.

Complex, tri- and tetra-antennary N-glycans are abundantly expressed in the gut, playing important roles in regulating membrane-bound and secreted proteins of the epithelium and mucosal T cell populations.^1, 31, 57, 58^ *Mgat4a/b* double-knockout animals (KO) do not make tri- and tetra-antennary N-glycans, but there is a compensatory up-regulation of downstream glycosyltransferases. As a result, *Mgat4a/b* double-KO animals develop longer polylactosamine (poly-LacNAc) extensions on bi-antennary glycan branches to compensate for loss of branching.^31^ These findings illustrate, “self-correction” mechanisms step in when N-glycan branching is impaired. Importantly, these “self-correction” processes are dependent on UDP-GlcNAc availability and the branching glycosyltransferases (e.g. MGAT1,2,4,5) and trans-Golgi extension enzymes (e.g. β1,3-N-acetylglucosaminyltransferases) that also require Mn.^30^ The tissue-specific effects of impaired “self-correction” and/or alterations in the length of polylactosamine extensions remain to be fully elucidated, yet likely alter glycoprotein signaling and stability at the plasma membrane, affect interaction with the galectin lattice, and influence host-microbiome interactions.^25, 59, 60^

The specific drivers of altered glycosyltransferase expression and the increase in truncated N-glycans in Crohn’s disease remain unclear, yet limited data implicate epithelial, immune and microbiota interactions.^61^ Aberrant bile acid homeostasis, decreased FGF19 signaling, and increased intestinal permeability are known contributors to intestinal inflammation and represent therapeutic targets in Crohn’s disease, increasing the possibility of clinical benefit with normalization via glycan-targeting therapy in carriers of ZIP8 391-Thr.^45, 62–67^ The effects of glycan-targeting therapy likely extend beyond the epithelium to the immune compartment where tri- and tetra-antennary N-glycans on T cells favor an immunoregulatory phenotype.^1, 40, 68^ We cannot exclude additional pathogenic mechanisms related to other cell-type specific activities of ZIP8 or perturbation of metal homeostasis beyond Mn, however integration of the observations made in humans linking genetic variation in ZIP8 to Mn homeostasis and glycosylation strongly prioritize aberrant N-glycosylation as a therapeutic target in carriers of ZIP8 391-Thr.^13, 69–72^

GlcNAc supplementation builds from prior studies in animal models and patients with IBD showing safety and therapeutic benefit, as well as more recent studies in the treatment of patients with multiple sclerosis, with immune-mediated effects of enhancing N-glycan branching on effector T cells.^1, 38–40, 73^ By lectin staining, physiologic measurements of bile acid homeostasis and intestinal permeability, and DSS challenge (tested only in Zip8^393T/393T^ mice), the effect of GlcNAc supplementation was most significant in Zip8^+/393T^ and Zip8^393T/393T^ mice with minimal effects on Zip8^+/+^ animals. These data support a specific benefit in ZIP8 391-Thr carriers, but the prior studies suggest that benefit from glycan-targeting therapy may extend to patients with other drivers of aberrant N-glycosylation and aligns with our data demonstrating consistent down-regulation of mRNA expression of glycosyltransferases and increased abundance of truncated N-glycans associated with Crohn’s disease compared to healthy individuals. These effects may be mediated through Mn availability. Notably, increased dietary Mn intake was recently identified as a protective against Crohn’s disease; while at the population-level, the intake of dietary Mn has decreased by >40% in the last 15 years.^74, 75^ Regulation of *de novo* UDP-GlcNAc availability is dependent on the hexosamine biosynthetic pathway that integrates the metabolism of carbohydrates, amino acids, fat, and nucleotides, with supply also determined by salvage pathways. Supply of GlcNAc in the gut may also be determined by resident microbiota and degradation of dietary-derived monosaccharides and glycopeptides.^76^ Together, the genetic associations between ZIP8 391-Thr and human pathology expose Mn homeostasis and aberrant N-glycosylation as contributors to disease pathogenesis. Multiple additional factors, like diet and metabolism, likely further modify disease risk in ZIP8 391-Thr carriers and may make others susceptible to dysfunction of these disease-associated pathways.

In summary, the common (global minor allele frequency of approximately 5%) and highly pleiotropic ZIP8 A391T has focused new attention on Mn homeostasis and Mn-dependent processes in human health and disease. Integration of human data from individuals with rare, loss-of-function mutations in ZIP8 directly link Mn status, N-glycosylation, and disease classification as a CDG. Treatment of defective glycosylation represents a novel target for adjunctive therapy in the care of patients with ZIP8 391-Thr, challenging the clinical paradigm that targeting defective glycosylation is reserved for patients with rare disease. Future clinical trials are needed to study the clinical benefit and underlying mechanisms of glycan-specific therapy across ZIP8 391-Thr associated disease states.

## Acknowledgments

The authors acknowledge the research contributions made by patients to allow for this research study. The authors acknowledge genotyping data made possible through the NIDDK IBD Genetics Consortium, coordinated by Lisa Datta; DK062431). Confocal imaging supported by the Johns Hopkins Conte Digestive Disease Core (P30-DK089502). All MALDI imaging experiments were performed in the Johns Hopkins Applied Imaging Mass Spectrometry (AIMS) Core. The authors acknowledge many helpful conversations with Drs. Mark Donowitz and Cynthia Sears (Johns Hopkins).

## Author Contribution

V.T.: conceptualization, methodology, analysis, visualization, writing, reviewing, editing. J.K.: analysis, visualization. R.L.: analysis and visualization. S.B.: resources, reviewing, editing. M.L.: resources, reviewing, editing. C.T.: analysis, visualization, writing, and reviewing. K.G.: analysis, visualization, and reviewing. N.Z.: analysis, visualization, and reviewing. J.M.: conceptualization, methodology, analysis, resources, writing the first draft of the manuscript, reviewing, editing, visualization, supervision, and funding acquisition.

## Materials and Methods

### Reagents

Dextran sodium sulfate (DSS; molecular weight 36-50 kDA, reagent grade) was purchased from MP Biochemicals LLC (Solon, OH), N-Acetylglucosamine USP supplement was purchased from Wellesley Therapeutics Inc.

### Human experimental design

Ileal biopsies embedded in formalin were obtained from Johns Hopkins Pathology for genotyped patients with Crohn’s disease. Patients were previously genotyped as part of participation in the NIDDK IBD Genetics Consortium. Additionally, ileal biopsies from healthy tissue was obtained from discarded and de-identified surgical tissue. The use of genetic data and human biospecimens was approved by the Johns Hopkins University School of Medicine institutional review board. Lectin immunofluorescence was performed on FFPE samples as outlined below in “immunostaining.”

A secondary analysis of publicly-available bulk RNAseq derived from human ileal biopsies was performed (GSE57945). A curated collection of 79 glycosyltransferase genes related to N-glycosylation were used in the analysis. Discovery determined using the two-stage linear step-up procedure of Benjamini, Krieger and Yekutieli, with Q = 1%. Each row was analyzed individually, without assuming a consistent SD. Number of t tests: 77 (not counting 2 row(s) where the t-test could not be computed). Volcano plot shown in Fig. 1 with analysis (Graphpad; **Supplementary Table 1**).

### Animals and experimental design

For the majority of our experiments, ten-twelve weeks old male wild-type (WT) C57BL/6 mice and Zip8 393T-KI mice were obtained from our colony unless noted. The mice were housed in a single facility in plastic cages, 4-5 mice per cage, at a constant temperature (25°C) and humidity (50 ±10%) and maintained in air-conditioned quarters with a 12-hour light/dark cycle. Mice were fed conventional chow (Harlan/Envigo Teklad Global 18% Protein Extruded Diet 2018SX) containing Mn 100 ppm with the exception of the animals used in the purified diet experiment controlling Mn content.

To study the effects of dietary Mn and host Mn status on ileal epithelial glycosylation, WT C57BL/6 male mice were purchased directly from Jackson Laboratory at 6 weeks of age. Upon arrival in our colony, the animals were fed purified diets containing variable Mn content (Inotiv Teklad TD.97184 with Mn <1 ppm or 2400 ppm). The purified diet differs from conventional chow both in Mn content as well as the carbohydrate source. Conventional chow consists of whole grains (corn, wheat, soybean meal), while the purified diet consists of casein, corn starch, sucrose, and cellulose. This is

For oral GlcNAc supplementation experiments, age and weight matched WT C57BL/6 mice and Zip8 393T-KI mice were co-housed and fed 0.25 mg/ml in drinking water for up to four weeks before sacrifice and tissue harvesting as outlined in figure legends.

For DSS experiments, mice of mixed genotypes were co-housed to mitigate cage-level effects WT C57BL/6 and Zip8 393T-KI mice (n=24) were co-housed 1 week prior to start of the experiment. Mice were raised under identical environmental conditions (e.g., 12 h light/dark cycle, identical food, and drinking water) and were randomly divided into three groups: (1) sham, (2) DSS alone, (3) DSS + GlcNAc 0.25 mg/ml. Sham animals were exposed only to distilled water. The GlcNAc groups were pre-treated with GlcNAc for 7 days prior to DSS exposure. To elicit colitis, mice were administered with 2% DSS in drinking water for 5 days. For the animals in the GlcNAc group, GlcNAc was maintained in the drinking water for the entirety of the experimental period, followed by administration with only distilled water for an additional 9 days (14 days after start of DSS). Changes in body weight and developmental of clinical symptoms were evaluated every day during the experimental period.

### Disease activity index (DAI) and histological observation

Colitis was evaluated using DAI, as described by Cooper et. al.^77^ with some modifications, using parameters of body-weight loss, stool consistency, and bleeding. Briefly, no weight loss was scored as 0 points, weight loss of 1-5%, 1 point; loss of 6-10%, 2 points; 11-20% weight loss as 3 points; and > 20% of weight loss as 4 points. Stool consistency was characterized as normal (0 point), loose (2 points), and diarrhea (4 points). No bleeding was scored as 0 point, mild bleeding as 2 points and gross bleeding as 4 points. All parameters were scored daily during DSS treatment. Scores assigned for each parameter were used to calculate the average DAI. Mice were sacrificed by isoflurane inhalation after the experimental period. Colon tissues were washed with PBS to remove feces and blood, and colon lengths were measured thereafter. Blood samples, feces and other tissues were stored at −80^°^C until assayed. The colon tissues and ileal tissues were preserved in 10% buffered formalin solutions and finally embedded in paraffin. After de-paraffinizing tissue sections (3 μm) on glass slides, the sections were stained with hematoxylin and eosin. Microscopic sections were examined, and histological review and scoring was performed; extent of inflammatory infiltrate, goblet cell loss, crypt density, crypt hyperplasia, muscle thickening, submucosal infiltrate, crypt abscesses, and ulceration were assessed.^78^

### Immunostaining

Paraformaldehyde-fixed and paraffin-embedded human and mouse ileal tissue sections were prepared on glass slides, deparaffinized with xylene, and rehydrated with a descending ethanol series into phosphate-buffered saline (PBS). Antigen retrieval was done by heat mediation in a citrate buffer (#S1699; Dako), and then sections were blocked with 4% serum for 1 h at room temperature. Sections were permeabilized with PBS containing 1% bovine serum albumin and 0.2% Triton X-100. After blocking, samples were incubated with Fluorescein dyes (PHA-L rhodamine conjugated lectin and succinylated WGA lectin) from Vector Labs (RL-1112 and FL-1021s respectively) and Hoechst at concentration of 10 µg/ml in blocking buffer for 1 hour at room temperature. After antibody staining and washes, slides were mounted on glass slides using Fluoro-gel (#17985-10; EMS) and cover slipped. Images were captured using a Zeiss LSM 700 confocal microscope (PHA-L 639, sWGA 488 and Hoechst 405) with a 20X objective. Controls, including GlcNAc competition and PNGase F incubation are included in **Supplementary Fig. 3**.

### Measurement of soluble CD14 levels

Soluble CD14 is a receptor for bacterial LPS with increased expression correlating with impaired intestinal barrier function.^43^ At terminal blood collection, blood will be processed for serum, and CD14 will be measured using a commercially available ELISA (R&D Systems) with optical density measured at 450 nm.

### Total bile acid measurements in mice

To study bile acid homeostasis, we measured total bile acid levels in the liver and stool. Briefly, mice were fasted overnight, and then stool was collected at the time of sacrifice. Approximately, 10 mg of liver tissues or feces was homogenized with 1 mL of cold isopropanol and heated at 50°C for 2h for extraction of bile acids. After bringing the extracts to room temperature, we centrifuged them at 2000g for 15 minutes and fecal or hepatic bile acid concentrations in the collected supernatant was measured using a commercial colorimetric total bile acid assay kit (Cell Biolabs # STA-631) as per manufacturer’s instructions. This assay is based on an enzyme driven reaction and the colorimetric kit requires the presence of NADH, and thio-NAD+. In the assay the bile acids are incubated in the presence of 3-alpha hydroxysteroiddehydrogenase (3α-HSD) and the reduction of thio-NAD+ to thio-NADH is detected by colorimetric absorbance. Each bile acid standard and samples was assayed in duplicate. A freshly prepared standard curve was used each time the assay was performed. Each sample replicate will require two paired wells, one to be treated with 3α-HSD and one without the enzyme (NADH). After incubation for 30-60 minutes at 37°C, the end point calorimetric results were read at a primary wavelength at 405 nm and a secondary wavelength 630 nm using a microplate spectrophotometer.

### mRNA measurements

RNA was extracted using the Direct-zol RNA Miniprep Kit (Zymo Research # R2052) as per the manufacturer’s protocols. Purified RNA concentrations were measured using wavelengths of 260/280 nm on the Beckman Coulter DU 800 Spectrophotometer. From equal amounts of RNA, cDNA was generated using iSCRIPT. Real-time quantitative PCR was performed and analyzed using Power SYBR Green Master Mix (Thermo Fisher Scientific) reagents on the QuantStudio 6 flex real time PCR system. The sequences of the sense and antisense primers were as follows: Cyp7a1 Fwd 5’-TCC CTC CTT TGA AAA ACG TG-3’, Cyp7a1 Rvs 5’-GAG GTT CTG AGG CTG TGC TC-3’; Fgf15 Fwd 5’-TGA GCC ATC CAG TTG TGT CC-3’, Fgf15 Rvs 5’-CCA CTG GAG AAT TTG GGG CT-3’, Gapdh Fwd 5’-CAT CAC TGC CAC CCA GAA GAC TG-3’. Gapdh Rvs 5’-ATG CCA GTG AGC TTC CCG TTC AG-3’. Gapdh was be included as an internal control, and relative expression was calculated using the 2(–delta delta Ct) method.

### MALDI Imaging

Sample preparation and MALDI imaging were performed as per Scupakova et al.^79^ Tissues were cut to 4 micron thickness onto washed and dried ITO slides (Delta Technologies, Loveland, CO). Samples were stored at room temperature prior to use. Slides were placed on a hot plate at 60 °C for 1 hour prior to PNGase F digestion. Paraffin was removed by washing in xylene (10 minutes) followed by rehydration in 100% ethanol (2 x 2 minutes) and HPLC grade water (2 x 5 minutes). Without drying, antigen retrieval was performed in a pressure cooker (TintoRetriever, Bio SB, Goleta, CA), on the low setting for 20 minutes in 10 mM citric acid buffer. After antigen retrieval slides were washed with HPLC grade water (2 x 1 minute). Slides were allowed to dry on the bench top. PNGase F (Bulldog Bio, Portsmouth, NH) was dissolved in 450 µL of HPLC grade water. PNGase F was sprayed using an HTX M3+ sprayer (HTX Technologies, Chapel Hill, NC) using the following method: nozzle temperature 30 °C, pressure 10 psi, flow rate 10 µL/min, velocity 900 mm/min, track spacing 2 mm, number of passes 15, pattern HH, and drying time 2 sec with a final linear flow rate of 1.11×10^−5^ mL/min. Slide was incubated in a humid chamber for 3 hours at 37 °C. After incubation, CHCA matrix (5 mg/mL in 50% acetonitrile with 0.2% TFA) was applied using an HTX M5 sprayer (HTX Technologies, Chapel Hill, NC) using the following method: nozzle temperature 30 °C, number of passes 8, flow rate 0.1 mL/min, velocity 750 mm/min, track spacing 2 mm, pattern CC, pressure 10 psi, and gas flow rate of 3. MALDI imaging was performed on a Bruker RapifleX ToF/ToF at 50 micron spatial resolution (50 micron raster with 50 micron single laser) in reflection positive mode with 200 laser shots per pixel at 10 kHz repetition rate. Data was imported into SCiLS lab and TIC normalized.

### Statistical analysis

Statistical analysis was performed using Prism Graphpad (RRID:SCR_000306). The data are expressed as the mean ± SEM. When two groups were compared, t-test was used with p <0.05 considered statistically significant. When three or more groups were compared, one-way and two-way ANOVA with multiple comparison testing were used with p <0.05 considered statistically significant. Additional details of statistical analyses and representation of statistical significance by asterisks included in figure legends.

### Human and animal study approvals

Human studies were approved by the Johns Hopkins University School of Medicine (JHU-SOM) Institutional Review Board (NA_00038329). All procedures involved animals were reviewed approved by the Animal Care and Use Committee at the Johns Hopkins University School of Medicine (MO22M349) in accordance with the Association for Assessment and Accreditation of Laboratory Animal Care International.

## References

1. Dias AM, Correia A, Pereira MS, Almeida CR, Alves I, Pinto V, Catarino TA, Mendes N, Leander M, Oliva-Teles MT, Maia L, Delerue-Matos C, Taniguchi N, Lima M, Pedroto I, Marcos-Pinto R, Lago P, Reis CA, Vilanova M, Pinho SS. Metabolic control of T cell immune response through glycans in inflammatory bowel disease. Proc Natl Acad Sci U S A. 2018;115(20):E4651–E60. doi: 10.1073/pnas.1720409115. PubMed PMID: 29720442; PMCID: PMC5960299.

2. Lin W, Vann DR, Doulias PT, Wang T, Landesberg G, Li X, Ricciotti E, Scalia R, He M, Hand NJ, Rader DJ. Hepatic metal ion transporter ZIP8 regulates manganese homeostasis and manganese-dependent enzyme activity. J Clin Invest. 2017;127(6):2407–17. doi: 10.1172/JCI90896. PubMed PMID: 28481222; PMCID: PMC5451243.

3. Choi EK, Rajendiran TM, Soni T, Park JH, Aring L, Muraleedharan CK, Garcia-Hernandez V, Kamada N, Samuelson LC, Nusrat A, Iwase S, Seo YA. The manganese transporter SLC39A8 links alkaline ceramidase 1 to inflammatory bowel disease. Nat Commun. 2024;15(1):4775. doi: 10.1038/s41467-024-49049-8. PubMed PMID: 38839750; PMCID: PMC11153611.

4. Pickrell JK, Berisa T, Liu JZ, Segurel L, Tung JY, Hinds DA. Detection and interpretation of shared genetic influences on 42 human traits. Nat Genet. 2016;48(7):709–17. doi: 10.1038/ng.3570. PubMed PMID: 27182965.

5. Schizophrenia Working Group of the Psychiatric Genomics C. Biological insights from 108 schizophrenia-associated genetic loci. Nature. 2014;511(7510):421–7. doi: 10.1038/nature13595. PubMed PMID: 25056061; PMCID: PMC4112379.

6. Speliotes EK, Willer CJ, Berndt SI, Monda KL, Thorleifsson G, Jackson AU, Lango Allen H, Lindgren CM, Luan J, Magi R, Randall JC, Vedantam S, Winkler TW, Qi L, Workalemahu T, Heid IM, Steinthorsdottir V, Stringham HM, Weedon MN, Wheeler E, Wood AR, Ferreira T, Weyant RJ, Segre AV, Estrada K, Liang L, Nemesh J, Park JH, Gustafsson S, Kilpelainen TO, Yang J, Bouatia-Naji N, Esko T, Feitosa MF, Kutalik Z, Mangino M, Raychaudhuri S, Scherag A, Smith AV, Welch R, Zhao JH, Aben KK, Absher DM, Amin N, Dixon AL, Fisher E, Glazer NL, Goddard ME, Heard-Costa NL, Hoesel V, Hottenga JJ, Johansson A, Johnson T, Ketkar S, Lamina C, Li S, Moffatt MF, Myers RH, Narisu N, Perry JR, Peters MJ, Preuss M, Ripatti S, Rivadeneira F, Sandholt C, Scott LJ, Timpson NJ, Tyrer JP, van Wingerden S, Watanabe RM, White CC, Wiklund F, Barlassina C, Chasman DI, Cooper MN, Jansson JO, Lawrence RW, Pellikka N, Prokopenko I, Shi J, Thiering E, Alavere H, Alibrandi MT, Almgren P, Arnold AM, Aspelund T, Atwood LD, Balkau B, Balmforth AJ, Bennett AJ, Ben-Shlomo Y, Bergman RN, Bergmann S, Biebermann H, Blakemore AI, Boes T, Bonnycastle LL, Bornstein SR, Brown MJ, Buchanan TA, Busonero F, Campbell H, Cappuccio FP, Cavalcanti-Proenca C, Chen YD, Chen CM, Chines PS, Clarke R, Coin L, Connell J, Day IN, den Heijer M, Duan J, Ebrahim S, Elliott P, Elosua R, Eiriksdottir G, Erdos MR, Eriksson JG, Facheris MF, Felix SB, Fischer-Posovszky P, Folsom AR, Friedrich N, Freimer NB, Fu M, Gaget S, Gejman PV, Geus EJ, Gieger C, Gjesing AP, Goel A, Goyette P, Grallert H, Grassler J, Greenawalt DM, Groves CJ, Gudnason V, Guiducci C, Hartikainen AL, Hassanali N, Hall AS, Havulinna AS, Hayward C, Heath AC, Hengstenberg C, Hicks AA, Hinney A, Hofman A, Homuth G, Hui J, Igl W, Iribarren C, Isomaa B, Jacobs KB, Jarick I, Jewell E, John U, Jorgensen T, Jousilahti P, Jula A, Kaakinen M, Kajantie E, Kaplan LM, Kathiresan S, Kettunen J, Kinnunen L, Knowles JW, Kolcic I, Konig IR, Koskinen S, Kovacs P, Kuusisto J, Kraft P, Kvaloy K, Laitinen J, Lantieri O, Lanzani C, Launer LJ, Lecoeur C, Lehtimaki T, Lettre G, Liu J, Lokki ML, Lorentzon M, Luben RN, Ludwig B, Magic, Manunta P, Marek D, Marre M, Martin NG, McArdle WL, McCarthy A, McKnight B, Meitinger T, Melander O, Meyre D, Midthjell K, Montgomery GW, Morken MA, Morris AP, Mulic R, Ngwa JS, Nelis M, Neville MJ, Nyholt DR, O’Donnell CJ, O’Rahilly S, Ong KK, Oostra B, Pare G, Parker AN, Perola M, Pichler I, Pietilainen KH, Platou CG, Polasek O, Pouta A, Rafelt S, Raitakari O, Rayner NW, Ridderstrale M, Rief W, Ruokonen A, Robertson NR, Rzehak P, Salomaa V, Sanders AR, Sandhu MS, Sanna S, Saramies J, Savolainen MJ, Scherag S, Schipf S, Schreiber S, Schunkert H, Silander K, Sinisalo J, Siscovick DS, Smit JH, Soranzo N, Sovio U, Stephens J, Surakka I, Swift AJ, Tammesoo ML, Tardif JC, Teder-Laving M, Teslovich TM, Thompson JR, Thomson B, Tonjes A, Tuomi T, van Meurs JB, van Ommen GJ, Vatin V, Viikari J, Visvikis-Siest S, Vitart V, Vogel CI, Voight BF, Waite LL, Wallaschofski H, Walters GB, Widen E, Wiegand S, Wild SH, Willemsen G, Witte DR, Witteman JC, Xu J, Zhang Q, Zgaga L, Ziegler A, Zitting P, Beilby JP, Farooqi IS, Hebebrand J, Huikuri HV, James AL, Kahonen M, Levinson DF, Macciardi F, Nieminen MS, Ohlsson C, Palmer LJ, Ridker PM, Stumvoll M, Beckmann JS, Boeing H, Boerwinkle E, Boomsma DI, Caulfield MJ, Chanock SJ, Collins FS, Cupples LA, Smith GD, Erdmann J, Froguel P, Gronberg H, Gyllensten U, Hall P, Hansen T, Harris TB, Hattersley AT, Hayes RB, Heinrich J, Hu FB, Hveem K, Illig T, Jarvelin MR, Kaprio J, Karpe F, Khaw KT, Kiemeney LA, Krude H, Laakso M, Lawlor DA, Metspalu A, Munroe PB, Ouwehand WH, Pedersen O, Penninx BW, Peters A, Pramstaller PP, Quertermous T, Reinehr T, Rissanen A, Rudan I, Samani NJ, Schwarz PE, Shuldiner AR, Spector TD, Tuomilehto J, Uda M, Uitterlinden A, Valle TT, Wabitsch M, Waeber G, Wareham NJ, Watkins H, Procardis C, Wilson JF, Wright AF, Zillikens MC, Chatterjee N, McCarroll SA, Purcell S, Schadt EE, Visscher PM, Assimes TL, Borecki IB, Deloukas P, Fox CS, Groop LC, Haritunians T, Hunter DJ, Kaplan RC, Mohlke KL, O’Connell JR, Peltonen L, Schlessinger D, Strachan DP, van Duijn CM, Wichmann HE, Frayling TM, Thorsteinsdottir U, Abecasis GR, Barroso I, Boehnke M, Stefansson K, North KE, McCarthy MI, Hirschhorn JN, Ingelsson E, Loos RJ. Association analyses of 249,796 individuals reveal 18 new loci associated with body mass index. Nat Genet. 2010;42(11):937–48. doi: 10.1038/ng.686. PubMed PMID: 20935630; PMCID: PMC3014648.

7. Haller G, McCall K, Jenkitkasemwong S, Sadler B, Antunes L, Nikolov M, Whittle J, Upshaw Z, Shin J, Baschal E, Cruchaga C, Harms M, Raggio C, Morcuende JA, Giampietro P, Miller NH, Wise C, Gray RS, Solnica-Krezel L, Knutson M, Dobbs MB, Gurnett CA. A missense variant in SLC39A8 is associated with severe idiopathic scoliosis. Nat Commun. 2018;9(1):4171. doi: 10.1038/s41467-018-06705-0. PubMed PMID: 30301978; PMCID: PMC6177404.

8. Nebert DW, Liu Z. SLC39A8 gene encoding a metal ion transporter: discovery and bench to bedside. Hum Genomics. 2019;13(Suppl 1):51. doi: 10.1186/s40246-019-0233-3. PubMed PMID: 31521203; PMCID: PMC6744627.

9. Initiative C-HG. A second update on mapping the human genetic architecture of COVID-19. Nature. 2023;621(7977):E7–E26. doi: 10.1038/s41586-023-06355-3. PubMed PMID: 37674002; PMCID: PMC10482689.

10. Bonaventura E, Barone R, Sturiale L, Pasquariello R, Alessandri MG, Pinto AM, Renieri A, Panteghini C, Garavaglia B, Cioni G, Battini R. Clinical, molecular and glycophenotype insights in SLC39A8-CDG. Orphanet J Rare Dis. 2021;16(1):307. doi: 10.1186/s13023-021-01941-y. PubMed PMID: 34246313; PMCID: PMC8272319.

11. Lairson LL, Henrissat B, Davies GJ, Withers SG. Glycosyltransferases: structures, functions, and mechanisms. Annu Rev Biochem. 2008;77:521–55. doi: 10.1146/annurev.biochem.76.061005.092322. PubMed PMID: 18518825.

12. Sladek V, Tvaroska I. First-Principles Interaction Analysis Assessment of the Manganese Cation in the Catalytic Activity of Glycosyltransferases. J Phys Chem B. 2017;121(25):6148–62. doi: 10.1021/acs.jpcb.7b03714. PubMed PMID: 28617600.

13. Tseng WC, Reinhart V, Lanz TA, Weber ML, Pang J, Le KXV, Bell RD, O’Donnell P, Buhl DL. Schizophrenia-associated SLC39A8 polymorphism is a loss-of-function allele altering glutamate receptor and innate immune signaling. Transl Psychiatry. 2021;11(1):136. doi: 10.1038/s41398-021-01262-5. PubMed PMID: 33608496; PMCID: PMC7895948 and owned and/or held options/restricted stock units for the company’s publicly traded shares.

14. Mealer RG, Jenkins BG, Chen CY, Daly MJ, Ge T, Lehoux S, Marquardt T, Palmer CD, Park JH, Parsons PJ, Sackstein R, Williams SE, Cummings RD, Scolnick EM, Smoller JW. The schizophrenia risk locus in SLC39A8 alters brain metal transport and plasma glycosylation. Sci Rep. 2020;10(1):13162. doi: 10.1038/s41598-020-70108-9. PubMed PMID: 32753748; PMCID: PMC7403432.

15. Ng E, Lind PM, Lindgren C, Ingelsson E, Mahajan A, Morris A, Lind L. Genome-wide association study of toxic metals and trace elements reveals novel associations. Hum Mol Genet. 2015;24(16):4739–45. doi: 10.1093/hmg/ddv190. PubMed PMID: 26025379; PMCID: PMC4512629.

16. Sunuwar L, Frkatovic A, Sharapov S, Wang Q, Neu HM, Wu X, Haritunians T, Wan F, Michel S, Wu S, Donowitz M, McGovern D, Lauc G, Sears C, Melia J. Pleiotropic ZIP8 A391T implicates abnormal manganese homeostasis in complex human disease. JCI Insight. 2020;5(20). doi: 10.1172/jci.insight.140978. PubMed PMID: 32897876; PMCID: PMC7605523.

17. Park JH, Hogrebe M, Gruneberg M, DuChesne I, von der Heiden AL, Reunert J, Schlingmann KP, Boycott KM, Beaulieu CL, Mhanni AA, Innes AM, Hortnagel K, Biskup S, Gleixner EM, Kurlemann G, Fiedler B, Omran H, Rutsch F, Wada Y, Tsiakas K, Santer R, Nebert DW, Rust S, Marquardt T. SLC39A8 Deficiency: A Disorder of Manganese Transport and Glycosylation. Am J Hum Genet. 2015;97(6):894–903. doi: 10.1016/j.ajhg.2015.11.003. PubMed PMID: 26637979; PMCID: PMC4678430.

18. Boycott KM, Beaulieu CL, Kernohan KD, Gebril OH, Mhanni A, Chudley AE, Redl D, Qin W, Hampson S, Kury S, Tetreault M, Puffenberger EG, Scott JN, Bezieau S, Reis A, Uebe S, Schumacher J, Hegele RA, McLeod DR, Galvez-Peralta M, Majewski J, Ramaekers VT, Care4Rare Canada C, Nebert DW, Innes AM, Parboosingh JS, Abou Jamra R. Autosomal-Recessive Intellectual Disability with Cerebellar Atrophy Syndrome Caused by Mutation of the Manganese and Zinc Transporter Gene SLC39A8. Am J Hum Genet. 2015;97(6):886–93. doi: 10.1016/j.ajhg.2015.11.002. PubMed PMID: 26637978; PMCID: PMC4678439.

19. Park JH, Mealer RG, Elias AF, Hoffmann S, Gruneberg M, Biskup S, Fobker M, Haven J, Mangels U, Reunert J, Rust S, Schoof J, Schwanke C, Smoller JW, Cummings RD, Marquardt T. N-glycome analysis detects dysglycosylation missed by conventional methods in SLC39A8 deficiency. J Inherit Metab Dis. 2020;43(6):1370–81. doi: 10.1002/jimd.12306. PubMed PMID: 32852845; PMCID: PMC8086894.

20. Mealer RG, Williams SE, Noel M, Yang B, D’Souza AK, Nakata T, Graham DB, Creasey EA, Cetinbas M, Sadreyev RI, Scolnick EM, Woo CM, Smoller JW, Xavier RJ, Cummings RD. The schizophrenia-associated variant in SLC39A8 alters protein glycosylation in the mouse brain. Mol Psychiatry. 2022;27(3):1405–15. doi: 10.1038/s41380-022-01490-1. PubMed PMID: 35260802; PMCID: PMC9106890.

21. Elliott LT, Sharp K, Alfaro-Almagro F, Shi S, Miller KL, Douaud G, Marchini J, Smith SM. Genome-wide association studies of brain imaging phenotypes in UK Biobank. Nature. 2018;562(7726):210-6. doi: 10.1038/s41586-018-0571-7. PubMed PMID: 30305740; PMCID: PMC6786974.

22. Hermann ER, Chambers E, Davis DN, Montgomery MR, Lin D, Chowanadisai W. Brain Magnetic Resonance Imaging Phenome-Wide Association Study With Metal Transporter Gene SLC39A8. Front Genet. 2021;12:647946. doi: 10.3389/fgene.2021.647946. PubMed PMID: 33790950; PMCID: PMC8005600.

23. Koropatkin NM, Cameron EA, Martens EC. How glycan metabolism shapes the human gut microbiota. Nat Rev Microbiol. 2012;10(5):323–35. doi: 10.1038/nrmicro2746. PubMed PMID: 22491358; PMCID: PMC4005082.

24. Varki A. Biological roles of glycans. Glycobiology. 2017;27(1):3–49. doi: 10.1093/glycob/cww086. PubMed PMID: 27558841; PMCID: PMC5884436.

25. Brazil JC, Parkos CA. Finding the sweet spot: glycosylation mediated regulation of intestinal inflammation. Mucosal Immunol. 2022;15(2):211–22. doi: 10.1038/s41385-021-00466-8. PubMed PMID: 34782709; PMCID: PMC8591159.

26. Li D, Achkar JP, Haritunians T, Jacobs JP, Hui KY, D’Amato M, Brand S, Radford-Smith G, Halfvarson J, Niess JH, Kugathasan S, Buning C, Schumm LP, Klei L, Ananthakrishnan A, Aumais G, Baidoo L, Dubinsky M, Fiocchi C, Glas J, Milgrom R, Proctor DD, Regueiro M, Simms LA, Stempak JM, Targan SR, Torkvist L, Sharma Y, Devlin B, Borneman J, Hakonarson H, Xavier RJ, Daly M, Brant SR, Rioux JD, Silverberg MS, Cho JH, Braun J, McGovern DP, Duerr RH. A pleiotropic missense variant in SLC39A8 is associated with Crohn’s disease and human gut microbiome composition. Gastroenterology. 2016;151(4):724–32. Epub 2016 Aug 1. doi: 10.1053/j.gastro.2016.06.051. PubMed PMID: 27492617; PMCID: PMC5037008.

27. Cummings RD EM. Antibodies and Lectins in Glycan Analysis. In: Varki A CR, Esko JD, et al., editor. Essentials of Glycobiology. Cold Spring Harbor (NY): Cold Spring Harbor Laboratory Press; 2009.

28. Haberman Y, Tickle TL, Dexheimer PJ, Kim MO, Tang D, Karns R, Baldassano RN, Noe JD, Rosh J, Markowitz J, Heyman MB, Griffiths AM, Crandall WV, Mack DR, Baker SS, Huttenhower C, Keljo DJ, Hyams JS, Kugathasan S, Walters TD, Aronow B, Xavier RJ, Gevers D, Denson LA. Pediatric Crohn disease patients exhibit specific ileal transcriptome and microbiome signature. J Clin Invest. 2014;124(8):3617–33. doi: 10.1172/JCI75436. PubMed PMID: 25003194; PMCID: PMC4109533.

29. Kugathasan S, Denson LA, Walters TD, Kim MO, Marigorta UM, Schirmer M, Mondal K, Liu C, Griffiths A, Noe JD, Crandall WV, Snapper S, Rabizadeh S, Rosh JR, Shapiro JM, Guthery S, Mack DR, Kellermayer R, Kappelman MD, Steiner S, Moulton DE, Keljo D, Cohen S, Oliva-Hemker M, Heyman MB, Otley AR, Baker SS, Evans JS, Kirschner BS, Patel AS, Ziring D, Trapnell BC, Sylvester FA, Stephens MC, Baldassano RN, Markowitz JF, Cho J, Xavier RJ, Huttenhower C, Aronow BJ, Gibson G, Hyams JS, Dubinsky MC. Prediction of complicated disease course for children newly diagnosed with Crohn’s disease: a multicentre inception cohort study. Lancet. 2017;389(10080):1710–8. doi: 10.1016/S0140-6736(17)30317-3. PubMed PMID: 28259484; PMCID: PMC5719489.

30. Mkhikian H, Mortales CL, Zhou RW, Khachikyan K, Wu G, Haslam SM, Kavarian P, Dell A, Demetriou M. Golgi self-correction generates bioequivalent glycans to preserve cellular homeostasis. Elife. 2016;5. doi: 10.7554/eLife.14814. PubMed PMID: 27269286; PMCID: PMC4940165.

31. Takamatsu S, Antonopoulos A, Ohtsubo K, Ditto D, Chiba Y, Le DT, Morris HR, Haslam SM, Dell A, Marth JD, Taniguchi N. Physiological and glycomic characterization of N-acetylglucosaminyltransferase-IVa and -IVb double deficient mice. Glycobiology. 2010;20(4):485–97. doi: 10.1093/glycob/cwp200. PubMed PMID: 20015870; PMCID: PMC2900882.

32. Ryczko MC, Pawling J, Chen R, Abdel Rahman AM, Yau K, Copeland JK, Zhang C, Surendra A, Guttman DS, Figeys D, Dennis JW. Metabolic Reprogramming by Hexosamine Biosynthetic and Golgi N-Glycan Branching Pathways. Sci Rep. 2016;6:23043. doi: 10.1038/srep23043. PubMed PMID: 26972830; PMCID: PMC4789752.

33. Schachter H. Biosynthetic controls that determine the branching and microheterogeneity of protein-bound oligosaccharides. Adv Exp Med Biol. 1986;205:53–85. doi: 10.1007/978-1-4684-5209-9_2. PubMed PMID: 3538817.

34. Grigorian A, Lee SU, Tian W, Chen IJ, Gao G, Mendelsohn R, Dennis JW, Demetriou M. Control of T Cell-mediated autoimmunity by metabolite flux to N-glycan biosynthesis. J Biol Chem. 2007;282(27):20027–35. doi: 10.1074/jbc.M701890200. PubMed PMID: 17488719.

35. Freeze HH. Perhaps a wee bit of sugar would help. Nat Genet. 2016;48(7):705–7. doi: 10.1038/ng.3600. PubMed PMID: 27350601; PMCID: PMC5428893.

36. Roth J, Ponzoni S, Aschner M. Manganese homeostasis and transport. Met Ions Life Sci. 2013;12:169–201. doi: 10.1007/978-94-007-5561-1_6. PubMed PMID: 23595673; PMCID: PMC6542352.

37. Peres TV, Schettinger MR, Chen P, Carvalho F, Avila DS, Bowman AB, Aschner M. “Manganese-induced neurotoxicity: a review of its behavioral consequences and neuroprotective strategies”. BMC Pharmacol Toxicol. 2016;17(1):57. doi: 10.1186/s40360-016-0099-0. PubMed PMID: 27814772; PMCID: PMC5097420.

38. Salvatore S, Heuschkel R, Tomlin S, Davies SE, Edwards S, Walker-Smith JA, French I, Murch SH. A pilot study of N-acetyl glucosamine, a nutritional substrate for glycosaminoglycan synthesis, in paediatric chronic inflammatory bowel disease. Aliment Pharmacol Ther. 2000;14(12):1567–79. doi: 10.1046/j.1365-2036.2000.00883.x. PubMed PMID: 11121904.

39. Zhu AP, I; Hidalgo, MP; Gandhi V. N-Acetylglucosamine for Treatment of Inflammatory Bowel Disease A real-world pragmatic clinical trial. Natural Medicine Journal. 2015;7(4):1–8.

40. Sy M, Newton BL, Pawling J, Hayama KL, Cordon A, Yu Z, Kuhle J, Dennis JW, Brandt AU, Demetriou M. N-acetylglucosamine inhibits inflammation and neurodegeneration markers in multiple sclerosis: a mechanistic trial. J Neuroinflammation. 2023;20(1):209. doi: 10.1186/s12974-023-02893-9. PubMed PMID: 37705084; PMCID: PMC10498575.

41. Muthusamy S, Malhotra P, Hosameddin M, Dudeja AK, Borthakur S, Saksena S, Gill RK, Dudeja PK, Alrefai WA. N-glycosylation is essential for ileal ASBT function and protection against proteases. Am J Physiol Cell Physiol. 2015;308(12):C964–71. doi: 10.1152/ajpcell.00023.2015. PubMed PMID: 25855079; PMCID: PMC4469748.

42. Briggs K, Tomar V, Ollberding N, Haberman Y, Bourgonje AR, Hu S, Chaaban L, Sunuwar L, Weersma RK, Denson LA, Melia JMP. Crohn’s Disease-Associated Pathogenic Mutation in the Manganese Transporter ZIP8 Shifts the Ileal and Rectal Mucosal Microbiota Implicating Aberrant Bile Acid Metabolism. Inflamm Bowel Dis. 2024. doi: 10.1093/ibd/izae003. PubMed PMID: 38289995.

43. Tabung FK, Birmann BM, Epstein MM, Martinez-Maza O, Breen EC, Wu K, Giovannucci EL. Influence of Dietary Patterns on Plasma Soluble CD14, a Surrogate Marker of Gut Barrier Dysfunction. Curr Dev Nutr. 2017;1(11). doi: 10.3945/cdn.117.001396. PubMed PMID: 29595830; PMCID: PMC5867900.

44. Nakata T, Creasey EA, Kadoki M, Lin H, Selig MK, Yao J, Lefkovith A, Daly MJ, Graham DB, Xavier RJ. A missense variant in SLC39A8 confers risk for Crohn’s disease by disrupting manganese homeostasis and intestinal barrier integrity. Proc Natl Acad Sci U S A. 2020;117(46):28930–8. doi: 10.1073/pnas.2014742117. PubMed PMID: 33139556; PMCID: PMC7682327.

45. Turpin W, Lee SH, Raygoza Garay JA, Madsen KL, Meddings JB, Bedrani L, Power N, Espin-Garcia O, Xu W, Smith MI, Griffiths AM, Moayyedi P, Turner D, Seidman EG, Steinhart AH, Marshall JK, Jacobson K, Mack D, Huynh H, Bernstein CN, Paterson AD, Crohn’s, Colitis Canada Genetic Environmental Microbial Project Research C, Abreu CGPrsdiM, Croitoru K. Increased Intestinal Permeability Is Associated With Later Development of Crohn’s Disease. Gastroenterology. 2020;159(6):2092–100 e5. doi: 10.1053/j.gastro.2020.08.005. PubMed PMID: 32791132.

46. Chassaing B, Rolhion N, de Vallee A, Salim SY, Prorok-Hamon M, Neut C, Campbell BJ, Soderholm JD, Hugot JP, Colombel JF, Darfeuille-Michaud A. Crohn disease--associated adherent-invasive E. coli bacteria target mouse and human Peyer’s patches via long polar fimbriae. J Clin Invest. 2011;121(3):966–75. doi: 10.1172/JCI44632. PubMed PMID: 21339647; PMCID: PMC3049390.

47. Pajusalu S, Vals MA, Mihkla L, Samarina U, Kahre T, Ounap K. The Estimated Prevalence of N-Linked Congenital Disorders of Glycosylation Across Various Populations Based on Allele Frequencies in General Population Databases. Front Genet. 2021;12:719437. doi: 10.3389/fgene.2021.719437. PubMed PMID: 34447415; PMCID: PMC8383291.

48. Chang IJ, He M, Lam CT. Congenital disorders of glycosylation. Ann Transl Med. 2018;6(24):477. doi: 10.21037/atm.2018.10.45. PubMed PMID: 30740408; PMCID: PMC6331365.

49. Wildeman AS, Patel NK, Cormack BP, Culotta VC. The role of manganese in morphogenesis and pathogenesis of the opportunistic fungal pathogen Candida albicans. PLoS Pathog. 2023;19(6):e1011478. doi: 10.1371/journal.ppat.1011478. PubMed PMID: 37363924; PMCID: PMC10328360.

50. Montllor-Albalate C, Kim H, Thompson AE, Jonke AP, Torres MP, Reddi AR. Sod1 integrates oxygen availability to redox regulate NADPH production and the thiol redoxome. Proc Natl Acad Sci U S A. 2022;119(1). doi: 10.1073/pnas.2023328119. PubMed PMID: 34969852; PMCID: PMC8740578.

51. Sazonovs A, Stevens CR, Venkataraman GR, Yuan K, Avila B, Abreu MT, Ahmad T, Allez M, Ananthakrishnan AN, Atzmon G, Baras A, Barrett JC, Barzilai N, Beaugerie L, Beecham A, Bernstein CN, Bitton A, Bokemeyer B, Chan A, Chung D, Cleynen I, Cosnes J, Cutler DJ, Daly A, Damas OM, Datta LW, Dawany N, Devoto M, Dodge S, Ellinghaus E, Fachal L, Farkkila M, Faubion W, Ferreira M, Franchimont D, Gabriel SB, Georges M, Gettler K, Giri M, Glaser B, Goerg S, Goyette P, Graham D, Hämäläinen E, Haritunians T, Heap GA, Hiltunen M, Hoeppner M, Horowitz JE, Irving P, Iyer V, Jalas C, Kelsen J, Khalili H, Kirschner BS, Kontula K, Koskela JT, Kugathasan S, Kupcinskas J, Lamb CA, Laudes M, Levine AP, Lewis J, Liefferinckx C, Loescher B-S, Louis E, Mansfield J, May S, McCauley JL, Mengesha E, Mni M, Moayyedi P, Moran CJ, Newberry R, O’Charoen S, Okou DT, Oldenburg B, Ostrer H, Palotie A, Pekow J, Peter I, Pierik MJ, Ponsioen CY, Pontikos N, Prescott N, Pulver AE, Rahmouni S, Rice DL, Saavalainen P, Sands B, Sartor RB, Schiff ER, Schreiber S, Schuum LP, Segal AW, Seksik P, Shawky R, Sheikh SZ, Silverberg M, Simmons A, Skeiceviciene J, Sokol H, Solomonson M, Somineni H, Sun D, Targan S, Turner D, Uhlig HH, van der Meulen AE, Vermeire S, Verstockt S, Voskuil MD, Winter HS, Young J, Consortium BI, IBD C-S, Consortium IIG, Consortium NIG, BioResource NI, Center RG, Consortium S, Network SI, Consortium UIG, Duerr RH, Franke A, Brant SR, Cho J, Weersma RK, Parkes M, Xavier R, Rivas MA, Rioux JD, McGovern D, Huang H, Anderson CA, Daly MJ. Sequencing of over 100,000 individuals identifies multiple genes and rare variants associated with Crohns disease susceptibility. medRxiv. 2021:2021.06.15.21258641. doi: 10.1101/2021.06.15.21258641.

52. Inflammatory Bowel Disease Exomes Portal [Website]. 2019 [cited 2019 August 23]. Available from: https://ibd.broadinstitute.org/.

53. Dulai PS, Singh S, Vande Casteele N, Boland BS, Rivera-Nieves J, Ernst PB, Eckmann L, Barrett KE, Chang JT, Sandborn WJ. Should We Divide Crohn’s Disease Into Ileum-Dominant and Isolated Colonic Diseases? Clin Gastroenterol Hepatol. 2019;17(13):2634–43. doi: 10.1016/j.cgh.2019.04.040. PubMed PMID: 31009791; PMCID: PMC6885453.

54. Cleynen I, Boucher G, Jostins L, Schumm LP, Zeissig S, Ahmad T, Andersen V, Andrews JM, Annese V, Brand S, Brant SR, Cho JH, Daly MJ, Dubinsky M, Duerr RH, Ferguson LR, Franke A, Gearry RB, Goyette P, Hakonarson H, Halfvarson J, Hov JR, Huang H, Kennedy NA, Kupcinskas L, Lawrance IC, Lee JC, Satsangi J, Schreiber S, Theatre E, van der Meulen-de Jong AE, Weersma RK, Wilson DC, International Inflammatory Bowel Disease Genetics C, Parkes M, Vermeire S, Rioux JD, Mansfield J, Silverberg MS, Radford-Smith G, McGovern DP, Barrett JC, Lees CW. Inherited determinants of Crohn’s disease and ulcerative colitis phenotypes: a genetic association study. Lancet. 2016;387(10014):156–67. doi: 10.1016/S0140-6736(15)00465-1. PubMed PMID: 26490195; PMCID: PMC4714968.

55. Atreya R, Siegmund B. Location is important: differentiation between ileal and colonic Crohn’s disease. Nat Rev Gastroenterol Hepatol. 2021;18(8):544–58. doi: 10.1038/s41575-021-00424-6. PubMed PMID: 33712743.

56. Magro F, Moreira PL, Catalano G, Alves C, Roseira J, Estevinho MM, Silva I, Dignass A, Peyrin-Biroulet L, Danese S, Jairath V, Dias CC, Santiago M. Has the therapeutical ceiling been reached in Crohn’s disease randomized controlled trials? A systematic review and meta-analysis. United European Gastroenterol J. 2023;11(2):202–17. doi: 10.1002/ueg2.12366. PubMed PMID: 36876515; PMCID: PMC10039796.

57. Wiggins CA, Munro S. Activity of the yeast MNN1 alpha-1,3-mannosyltransferase requires a motif conserved in many other families of glycosyltransferases. Proc Natl Acad Sci U S A. 1998;95(14):7945–50. doi: 10.1073/pnas.95.14.7945. PubMed PMID: 9653120; PMCID: PMC20909.

58. Nagae M, Yamaguchi Y, Taniguchi N, Kizuka Y. 3D Structure and Function of Glycosyltransferases Involved in N-glycan Maturation. Int J Mol Sci. 2020;21(2). doi: 10.3390/ijms21020437. PubMed PMID: 31936666; PMCID: PMC7014118.

59. Lau KS, Partridge EA, Grigorian A, Silvescu CI, Reinhold VN, Demetriou M, Dennis JW. Complex N-glycan number and degree of branching cooperate to regulate cell proliferation and differentiation. Cell. 2007;129(1):123–34. doi: 10.1016/j.cell.2007.01.049. PubMed PMID: 17418791.

60. Nabi IR, Shankar J, Dennis JW. The galectin lattice at a glance. J Cell Sci. 2015;128(13):2213–9. doi: 10.1242/jcs.151159. PubMed PMID: 26092931.

61. Nagao-Kitamoto H, Leslie JL, Kitamoto S, Jin C, Thomsson KA, Gillilland MG, 3rd, Kuffa P, Goto Y, Jenq RR, Ishii C, Hirayama A, Seekatz AM, Martens EC, Eaton KA, Kao JY, Fukuda S, Higgins PDR, Karlsson NG, Young VB, Kamada N. Interleukin-22-mediated host glycosylation prevents Clostridioides difficile infection by modulating the metabolic activity of the gut microbiota. Nat Med. 2020. doi: 10.1038/s41591-020-0764-0. PubMed PMID: 32066975.

62. Franzosa EA, Sirota-Madi A, Avila-Pacheco J, Fornelos N, Haiser HJ, Reinker S, Vatanen T, Hall AB, Mallick H, McIver LJ, Sauk JS, Wilson RG, Stevens BW, Scott JM, Pierce K, Deik AA, Bullock K, Imhann F, Porter JA, Zhernakova A, Fu J, Weersma RK, Wijmenga C, Clish CB, Vlamakis H, Huttenhower C, Xavier RJ. Gut microbiome structure and metabolic activity in inflammatory bowel disease. Nat Microbiol. 2019;4(2):293–305. doi: 10.1038/s41564-018-0306-4. PubMed PMID: 30531976; PMCID: PMC6342642.

63. Martini GR, Tikhonova E, Rosati E, DeCelie MB, Sievers LK, Tran F, Lessing M, Bergfeld A, Hinz S, Nikolaus S, Kumpers J, Matysiak A, Hofmann P, Saggau C, Schneiders S, Kamps AK, Jacobs G, Lieb W, Maul J, Siegmund B, Seegers B, Hinrichsen H, Oberg HH, Wesch D, Bereswill S, Heimesaat MM, Rupp J, Kniemeyer O, Brakhage AA, Brunke S, Hube B, Aden K, Franke A, Iliev ID, Scheffold A, Schreiber S, Bacher P. Selection of cross-reactive T cells by commensal and food-derived yeasts drives cytotoxic T(H)1 cell responses in Crohn’s disease. Nat Med. 2023;29(10):2602–14. doi: 10.1038/s41591-023-02556-5. PubMed PMID: 37749331; PMCID: PMC10579100.

64. Sinha SR, Haileselassie Y, Nguyen LP, Tropini C, Wang M, Becker LS, Sim D, Jarr K, Spear ET, Singh G, Namkoong H, Bittinger K, Fischbach MA, Sonnenburg JL, Habtezion A. Dysbiosis-Induced Secondary Bile Acid Deficiency Promotes Intestinal Inflammation. Cell Host Microbe. 2020;27(4):659–70 e5. doi: 10.1016/j.chom.2020.01.021. PubMed PMID: 32101703; PMCID: PMC8172352.

65. Thomas JP, Modos D, Rushbrook SM, Powell N, Korcsmaros T. The Emerging Role of Bile Acids in the Pathogenesis of Inflammatory Bowel Disease. Front Immunol. 2022;13:829525. doi: 10.3389/fimmu.2022.829525. PubMed PMID: 35185922; PMCID: PMC8850271.

66. Gadaleta RM, Garcia-Irigoyen O, Cariello M, Scialpi N, Peres C, Vetrano S, Fiorino G, Danese S, Ko B, Luo J, Porru E, Roda A, Sabba C, Moschetta A. Fibroblast Growth Factor 19 modulates intestinal microbiota and inflammation in presence of Farnesoid X Receptor. EBioMedicine. 2020;54:102719. doi: 10.1016/j.ebiom.2020.102719. PubMed PMID: 32259714; PMCID: PMC7136604.

67. Ahmad R, Sorrell MF, Batra SK, Dhawan P, Singh AB. Gut permeability and mucosal inflammation: bad, good or context dependent. Mucosal Immunol. 2017;10(2):307–17. doi: 10.1038/mi.2016.128. PubMed PMID: 28120842; PMCID: PMC6171348.

68. Brandt AU, Sy M, Bellmann-Strobl J, Newton BL, Pawling J, Zimmermann HG, Yu Z, Chien C, Dorr J, Wuerfel JT, Dennis JW, Paul F, Demetriou M. Association of a Marker of N-Acetylglucosamine With Progressive Multiple Sclerosis and Neurodegeneration. JAMA Neurol. 2021;78(7):842–52. doi: 10.1001/jamaneurol.2021.1116. PubMed PMID: 33970182; PMCID: PMC8111565 personal fees from Guthy Jackson Foundation, Einstein Foundation, BMB, and Deutsche Forschungsgemeinschaft Exc 157 during the conduct of the study; being cofounder and receiving shares from Motognosis GmbH and Nocturne GmbH outside the submitted work; and having a patent for GlcNAc as Serum Biomarker for Multiple Sclerosis issued. Dr Sy reported having an ownership stake in Glixis Therapeutics LLC outside the submitted work. Dr Bellmann-Strobl reported receiving nonfinancial support from Biogen Idec; personal fees from Bayer Healthcare, Merck Serono, Teva Pharmaceuticals, Roche, and Novartis; and personal fees and nonfinancial support from Sanofi Genzyme outside the submitted work. Dr Zimmermann reported receiving grants from Novartis and personal fees from Bayer Healthcare outside the submitted work. Ms Chien reported receiving speaking fees from Bayer and research funding from Novartis unrelated to this study. Dr Dorr reported receiving grants from Bayer during the conduct of the study; and personal fees from Bayer, Biogen, Novartis, Roche, Merck Serono, Sanofi, and Teva outside the submitted work. Dr Wuerfel reported receiving grants from the EU (Horizon2020) and the Swiss National Science Foundation; and serving on advisory boards for Actelion, Genzyme-Sanofi, Idorsia, InmuneBio, Novartis, and Roche. Dr Dennis reported holding a patent pending for PCT/US16/15807 N-Acetyl Glucosamine as a Biomarker of MS Disease Course and being an inventor on this patent, a patent on methods and compositions for preventing and treating a disease related to glycan dysregulation issued to Wellsley Therapeutics, and a patent on Analogs of N-acetylglucosamine pending; and being a cofounder of and holding shares in Glixis Therapeutics LLC. Dr Paul reported receiving grants from Celgene, Novartis, BMBF, Alexion, Guthy Jackson Foundation, Falck-Foundation, Roche, Almirall, Deutsche Forschungsgemeinschaft, Einstein Foundation, Biogen, and Merck Serono; research support and personal fees from UCB, Roche, Alexion, Sanofi Genzyme, Mitsubishi Tanabe, Bayer, Merck Serono, and Viela Bio outside the submitted work. Dr Demetriou reported receiving grants from the National Institute of Allergy and Infectious Disease and the National Center for Complementary and Integrative Health during the conduct of the study; having a patent for US9775859B2 issued, a patent for US10495646B2 issued, and a patent pending for US20170042919A1; and being a cofounder and shareholder of Glixis Therapeutics. No other disclosures were reported.

69. Liu MJ, Bao S, Galvez-Peralta M, Pyle CJ, Rudawsky AC, Pavlovicz RE, Killilea DW, Li C, Nebert DW, Wewers MD, Knoell DL. ZIP8 regulates host defense through zinc-mediated inhibition of NF-kappaB. Cell Rep. 2013;3(2):386–400. doi: 10.1016/j.celrep.2013.01.009. PubMed PMID: 23403290; PMCID: PMC3615478.

70. Samuelson DR, Haq S, Knoell DL. Divalent Metal Uptake and the Role of ZIP8 in Host Defense Against Pathogens. Front Cell Dev Biol. 2022;10:924820. doi: 10.3389/fcell.2022.924820. PubMed PMID: 35832795; PMCID: PMC9273032.

71. Sunuwar L, Tomar V, Wildeman A, Culotta V, Melia J. Hepatobiliary manganese homeostasis is dynamic in the setting of inflammation or infection in mice. FASEB J. 2023;37(9):e23123. doi: 10.1096/fj.202300539R. PubMed PMID: 37561548.

72. Winslow JWW, Limesand KH, Zhao N. The Functions of ZIP8, ZIP14, and ZnT10 in the Regulation of Systemic Manganese Homeostasis. Int J Mol Sci. 2020;21(9). doi: 10.3390/ijms21093304. PubMed PMID: 32392784; PMCID: PMC7246657.

73. Choi SI, Shin YC, Lee JS, Yoon YC, Kim JM, Sung MK. N-Acetylglucosamine and its dimer ameliorate inflammation in murine colitis by strengthening the gut barrier function. Food Funct. 2023;14(18):8533–44. doi: 10.1039/d3fo00282a. PubMed PMID: 37655824.

74. Choi EK, Aring L, Das NK, Solanki S, Inohara N, Iwase S, Samuelson LC, Shah YM, Seo YA. Impact of dietary manganese on experimental colitis in mice. FASEB J. 2020;34(2):2929–43. doi: 10.1096/fj.201902396R. PubMed PMID: 31908045; PMCID: PMC8103308.

75. Braun T, Feng R, Amir A, Levhar N, Shacham H, Mao R, Hadar R, Toren I, Algavi Y, Abu-Saad K, Zhuo S, Efroni G, Malik A, Picard O, Yavzori M, Agranovich B, Liu TC, Stappenbeck TS, Denson L, Kalter-Leibovici O, Gottlieb E, Borenstein E, Elinav E, Chen M, Ben-Horin S, Haberman Y. Diet-omics in the Study of Urban and Rural Crohn disease Evolution (SOURCE) cohort. Nat Commun. 2024;15(1):3764. doi: 10.1038/s41467-024-48106-6. PubMed PMID: 38704361.

76. Crouch LI, Urbanowicz PA, Basle A, Cai ZP, Liu L, Voglmeir J, Melo Diaz JM, Benedict ST, Spencer DIR, Bolam DN. Plant N-glycan breakdown by human gut Bacteroides. Proc Natl Acad Sci U S A. 2022;119(39):e2208168119. doi: 10.1073/pnas.2208168119. PubMed PMID: 36122227; PMCID: PMC9522356.

77. Cooper HS, Murthy SN, Shah RS, Sedergran DJ. Clinicopathologic study of dextran sulfate sodium experimental murine colitis. Lab Invest. 1993;69(2):238–49. PubMed PMID: 8350599.

78. Koelink PJ, Wildenberg ME, Stitt LW, Feagan BG, Koldijk M, van’t Wout AB, Atreya R, Vieth M, Brandse JF, Duijst S, Te Velde AA, D’Haens G, Levesque BG, van den Brink GR. Development of Reliable, Valid and Responsive Scoring Systems for Endoscopy and Histology in Animal Models for Inflammatory Bowel Disease. J Crohns Colitis. 2018;12(7):794–803. doi: 10.1093/ecco-jcc/jjy035. PubMed PMID: 29608662; PMCID: PMC6022651.

79. Scupakova K, Adelaja OT, Balluff B, Ayyappan V, Tressler CM, Jenkinson NM, Claes BS, Bowman AP, Cimino-Mathews AM, White MJ, Argani P, Heeren RM, Glunde K. Clinical importance of high-mannose, fucosylated, and complex N-glycans in breast cancer metastasis. JCI Insight. 2021;6(24). doi: 10.1172/jci.insight.146945. PubMed PMID: 34752419; PMCID: PMC8783675.

