## Supplementary Figures for "Aberrant N-glycosylation is a therapeutic target in carriers of a common and highly pleiotropic mutation in the manganese transporter ZIP8"

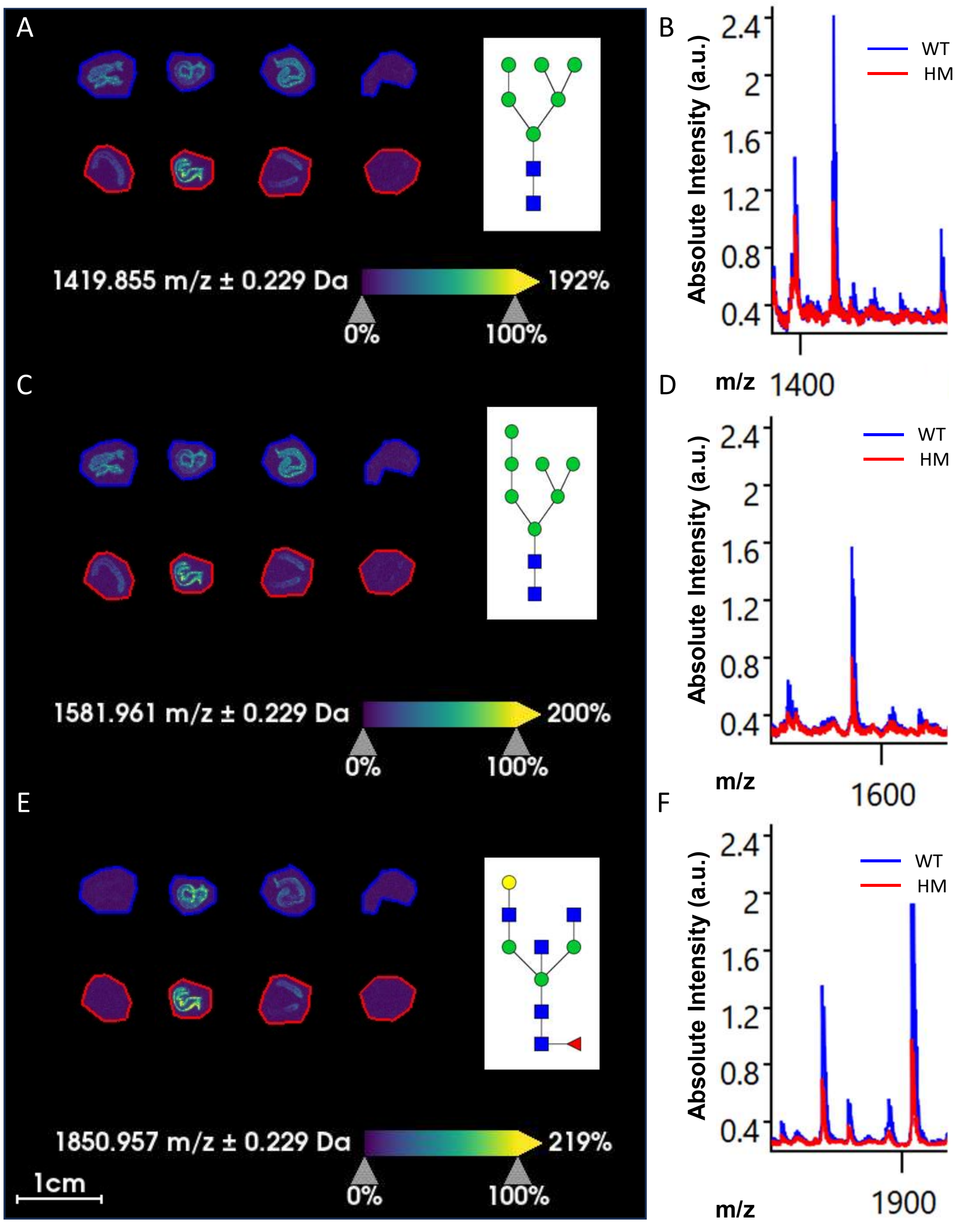


**Supplementary Figure 1.** Representative known glycan species from matrix-associated laser desorption/ionization (MALDI) mass spectrometry imaging (MSI). Glycan species identified as per Scupakova et al. Glycan representation as per convention (Fig. 1) of species at 1419.855 m/z (A,B), 1581.961 m/z (C,D), 1850.957 m/z (E,F).

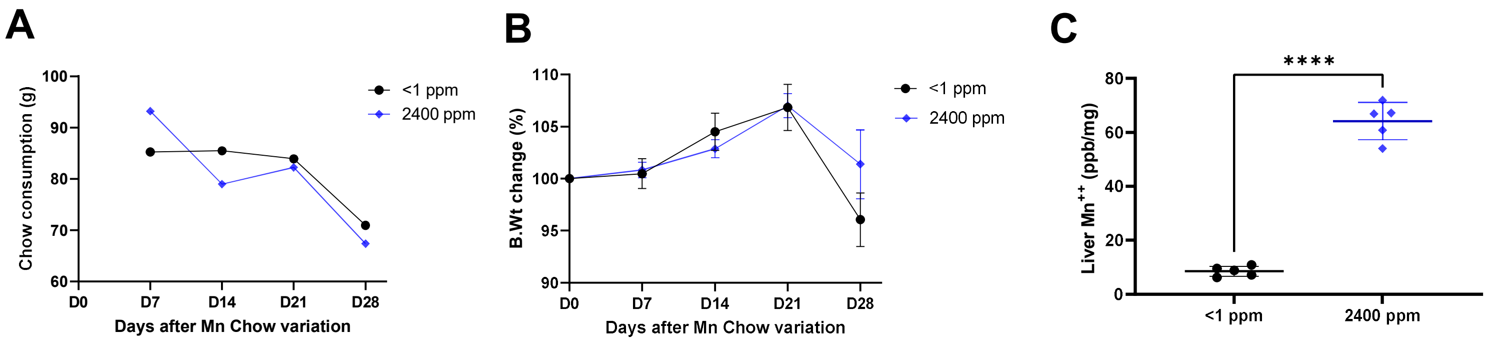


**Supplementary Figure 2. Variable dietary Mn feeding models Mn deficiency and excess.** C57BL/6 mice from Jackson Lab were fed purified diets containing <1 ppm or 2400 ppm Mn for 4 weeks from age 6 weeks to age 10 weeks. N = 5 male mice/group. **(A)** Chow consumption and **(B)** Body weight percent change did not differ between the two groups. Note the body weight measured at Day 28 (D28) was measured after an overnight fast prior to sacrifice, accounting for the observed decrease. **(C)** Total liver Mn, reflective of host Mn status, showed significant differences between the two diets. AAS. Statistical significance determined by two-sided t-test with four asterisks indicating $p < 0.00001$.

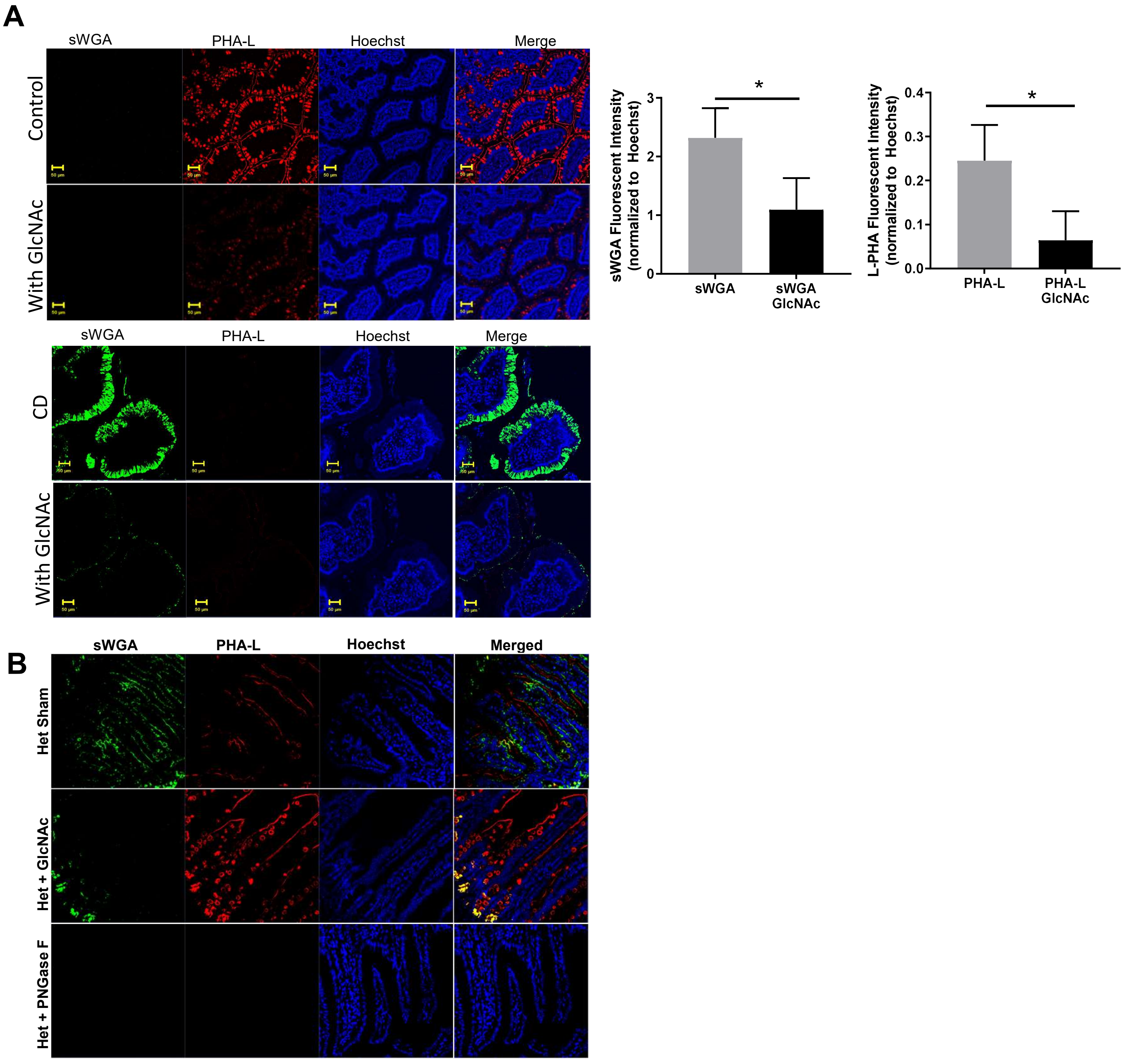


**Supplementary Figure 3.** Controls for lectin immunofluorescence. **(A)** Inhibition of Immunofluorescence of lectins in Healthy Controls and CD ileal human tissues by GlcNAc: Confocal laser-scanning triple-label immunofluorescence microscopy images of ileal human tissues paraffin sections, stained for PHA-L (red), sWGA (green), Hoechst (blue) and merged. 200 mM GlcNAc was incubated with Fluorescein dyes (PHA-L 639, sWGA 488 and Hoechst 405) 10 µg/ml in blocking buffer for 30-60 minutes at room temperature. After that samples were incubated with inhibited dyes for 1 hour at room temperature. Scale bar: 50um. **(B)** Demonstration of PNGase F effect. On-slide PNGase F digestion performed, abrogating sWGA and PHA-L staining.
